# Temporal single cell transcriptome atlas of zebrafish anterior segment development reveals high degree of conservation between the trabecular meshwork and the annular ligament

**DOI:** 10.1101/2022.10.22.513353

**Authors:** Oliver Vöcking, J.K. Famulski

## Abstract

Anterior segment dysgenesis (ASD), resulting in vision impairment, stems from maldevelopment of anterior segment (AS) tissues. Incidence of ASD has been linked to malfunction of periocular mesenchyme cells (POM). POM cells specify into anterior segment mesenchyme (ASM) cells which colonize and produce AS tissues. In this study we uncover ASM developmental trajectories associated with formation of the AS. Using a transgenic line of zebrafish that fluorescently labels the ASM throughout development, Tg[*foxc1b*:GFP], we isolated GFP+ ASM cells at several developmental timepoints (48-144hpf) and performed single cell RNA sequencing. Clustering analysis indicates subdifferentiation of ASM as early as 48hpf and subsequent diversification into corneal, epithelium/endothelium/stroma, or annular ligament (AL) lineages. Tracking individual clusters reveals common developmental pathways, up to 72hpf, for the AL and corneal endothelium/stroma, and distinct pathways for corneal epithelium starting at 48hpf. Spatiotemporal validation of over 80 genes found associated AS development demonstrates high degree of conservation with mammalian trabecular meshwork and corneal tissues. In addition, we characterize thirteen novel genes associated with AL and seven with corneal development. Overall, the data provide a molecular verification of the long-standing hypothesis that POM derived ASM give rise to AS tissues and highlight the high degree of conservation between zebrafish and mammals.

## INTRODUCTION

The development of the eye is a complex process that includes the orchestration of tissue types from different origins. One of these tissue types is the periocular mesenchyme, a subgroup of neural crest cells, that gives rise to the tissues of the anterior segment (AS) of the eye. Two major components of the anterior segment are the cornea and iridocorneal angle (ICA). The former’s main task is to focus incoming light onto the retina, while the latter is critically involved in regulating the intraocular pressure of the eye (1–6). Maldevelopment of the anterior segment and particularly these two components, can have severe consequences for eye function and can lead to blindness in the form of cataracts or glaucoma (4, 7).

Amongst several well-established model organisms used in the study of AS development, zebrafish embryos offer a unique opportunity to not only visualize AS development in real time but also quarry its components at numerous intervals and in high quantity. The mature zebrafish cornea is organized in five layers like the mammalian cornea: (1) the epithelium, (2) Bowman’s layer, (3) stroma, (4) Descement’s layer and (5) the endothelium (8–10). The corneal epithelium connects the cornea with the outside world. Underneath the epithelium and anterior to the stroma is the Bowman’s layer, which mainly consists of collagens (Collagen IV and VII). This is followed by the actual stroma (stroma proper), which is mainly made of extracellular matrix proteins, for the most part specifically organized collagens (mainly collagen I). The stroma itself is acellular but contains keratinocytes, which originate from neural crest cells. This is followed by the acellular Descement’s layer which is the basement membrane of the monolayered corneal endothelium (8, 10, 11). The main differences between the mammalian and zebrafish cornea are, that the corneal epithelium of zebrafish is thicker (circa 60% of the entire corneal thickness), likely due to the absence of eyelids and immediate exposure to the surrounding water medium. Also, in mammals the corneal stroma is the main contributor to the corneal thickness, whereas it makes up only approximately 30% of the thickness in zebrafish (10). The initial development of the zebrafish cornea appears very rapid. At 30hpf, a two-layered epithelium and a rudimentary stroma are already present. However, further differentiation and maturation is a much slower process. For instance, it takes 5 days after fertilization for the Bowman’s layer to appear, and keratinocytes only start invading the stroma between 14-28 dpf. Full maturity of the cornea is reached after approximately two months (10). The cornea has been a matter of research interest for some time and several marker genes are known, whose mutation can cause maldevelopments (3, 12). However, a detailed analysis of gene expression during early development is largely missing.

In contrast to the cornea, the iridocorneal angle (ICA) of zebrafish is significantly less studied. In mammals, the main parts of the ICA are the trabecular meshwork (TM) and the Schlemm’s canal (SC). These tissues have important roles in regulating the flow and drainage of the aqueous humor of the eye and thereby maintaining proper intraocular pressure (IOP) (4, 7, 13). In the absence of proper flow and drainage, the IOP can increase which in turn can lead to glaucoma, one of the leading causes of blindness in the US and worldwide (4, 7, 13–15). Accordingly, an increasing amount of attention has been paid to understand the trabecular meshwork (TM) and the Schlemm’s canal (SC), specifically the underlying genetics (4, 16–19). These studies catalogued gene expression from adult tissues at the singe cell level and in doing so, uncovered numerous potential targets for the study of ICA function. A potential roadblock to taking advantage of this treasure trove of data is the laborious nature of ICA functional examination in current models such as mice or primates. Zebrafish have become a well-accepted model for ocular disease modeling, in particular for retinopathies, however their use in ASD research has been less pronounced (8, 20–25). One potential issue is the physiological difference between mammalian and zebrafish ICA. The tissue within the zebrafish ICA is called the annular ligament (AL), which is similarly organized to the TM and SC of mammals and fulfills the same function (8, 9, 26, 27). It is a porous tissue, filled with glycoproteins and leads to a network of channels ending in the episcleral vasculature, like the SC. Its function is to regulate flow and drainage of the aqueous humor and thereby the intraocular pressure. In contrast to mammalian ICA with drains circumferentially, the AL facilitates drainage dorsally and aqueous humor production ventrally. Furthermore, certain previous studies suggested that the AL may not be a reliable a model for the human TM due to differences in the protein tested to reside in the tissues (26). However, at present we know very little about the actual genetic and molecular profile of the AL and a detailed characterization would help to determine it’s usefulness as a model for human TM diseases like glaucoma (28).

While recent studies have revealed some of the molecular composition of the adult cornea and ICA, there is still a lot more to uncover, particularly regarding the molecular mechanism regulating their development (16–19, 29–32). Work from various model systems, including zebrafish, chick, mouse, and human tissue, has firmly established the physiological landscape and developmental timing for the assembly of the AS (3, 29, 33–37). It has also affirmed the conservation and physiological function between such distinct species. The missing piece of the puzzle is how each of the tissues comprising the AS becomes functional and how this is regulated during development. Having a molecular atlas of these critical developmental events would go a long way to aid in the detection and possibly prevention of congenital blinding disorders associated with malformation of the AS. Novel approaches and advancements in the field of transcriptomics give an opportunity to finally begin such investigations. Recently, there has been an explosion of single cell transcriptomic analysis pertaining to retinal development, however, data for the development of the AS and particularly the TM are still sparse (17–19, 29). Furthermore, the current single cell transcriptomic data of the vertebrate eye includes tissue from a single, mostly adult time point. This includes recent work on human as well as monkey and pig AS samples (16–19, 30–32). As such, in our current study we used developing zebrafish to systematically analyze transcriptomic changes within ASM isolated from the AS over the course of early development. From our analysis of single cell transcriptomes, we have established clear developmental trajectories for progenitor cells of the cornea and annular ligament while uncovering new biological pathways that may be involved in the development of the zebrafish AS. Furthermore, we highlight that the annular ligament, while discounted in previous studies as divergent from the mammalian drainage structures, does exhibit gene expression patterns conserved with those of the trabecular meshwork (8, 9, 26). Overall, our study yields the first developmental atlas of AS development with single cell resolution and further supports that zebrafish can be used as model organism to study human diseases affecting the cornea and iridocorneal angle.

## MATERIALS AND METHODS

### Zebrafish maintenance

Zebrafish were maintained and treated according to the IACUC standards of the University of Kentucky (protocol # 2021-3781). The embryos were initially raised in embryo media (E3) at 28°C for 24 hours and thereafter in E3 medium containing 1-phenyl 2-thiourea (PTU) to maintain embryo transparency. The transgenic line Tg[*foxc1b*:GFP] was a gift from Dr. Brian Link. As wildtype we used the AB line.

### Whole-Mount in situ hybridization (WISH)

Whole mount in situ hybridizations were performed as previously described (38). Briefly, larvae of different developmental stages (24hpf, 3dpf and 5dpf) were fixed overnight in 4%PFA in PBS. Antisense RNA probes were generated by means of PCR, including T7 promoters in the primers and subsequent transcription with T7 Polymerase (Roche). Primer sequences used are found in table 1. Images of the embryos were taken with a Nikon Digital Sight DS-U3 camera and Elements software. Image adjustments for brightness and contrast were done with Adobe Photoshop and figures were assembled using Adobe Illustrator. 5dpf embryos, grown without PTU, were euthanized using MS222 and subsequently fixed in 4% paraformaldehyde for 24 hours. Fixed embryos were treated in 10% and subsequently 30% sucrose for 24hours each. Post sucrose washes, the embryos were mounted in OCT (optimal cutting temperature) embedding medium (Sakura) and frozen in cryosection molds. Cryosections were collected at 10μm thickness using a Leica cryotome. Images were collected using a Nikon Ti2 compound microscope with a 20X NA 0.95 objective. Subsequent processing was performed using Adobe photoshop and illustrator.

**TABLE 1.**
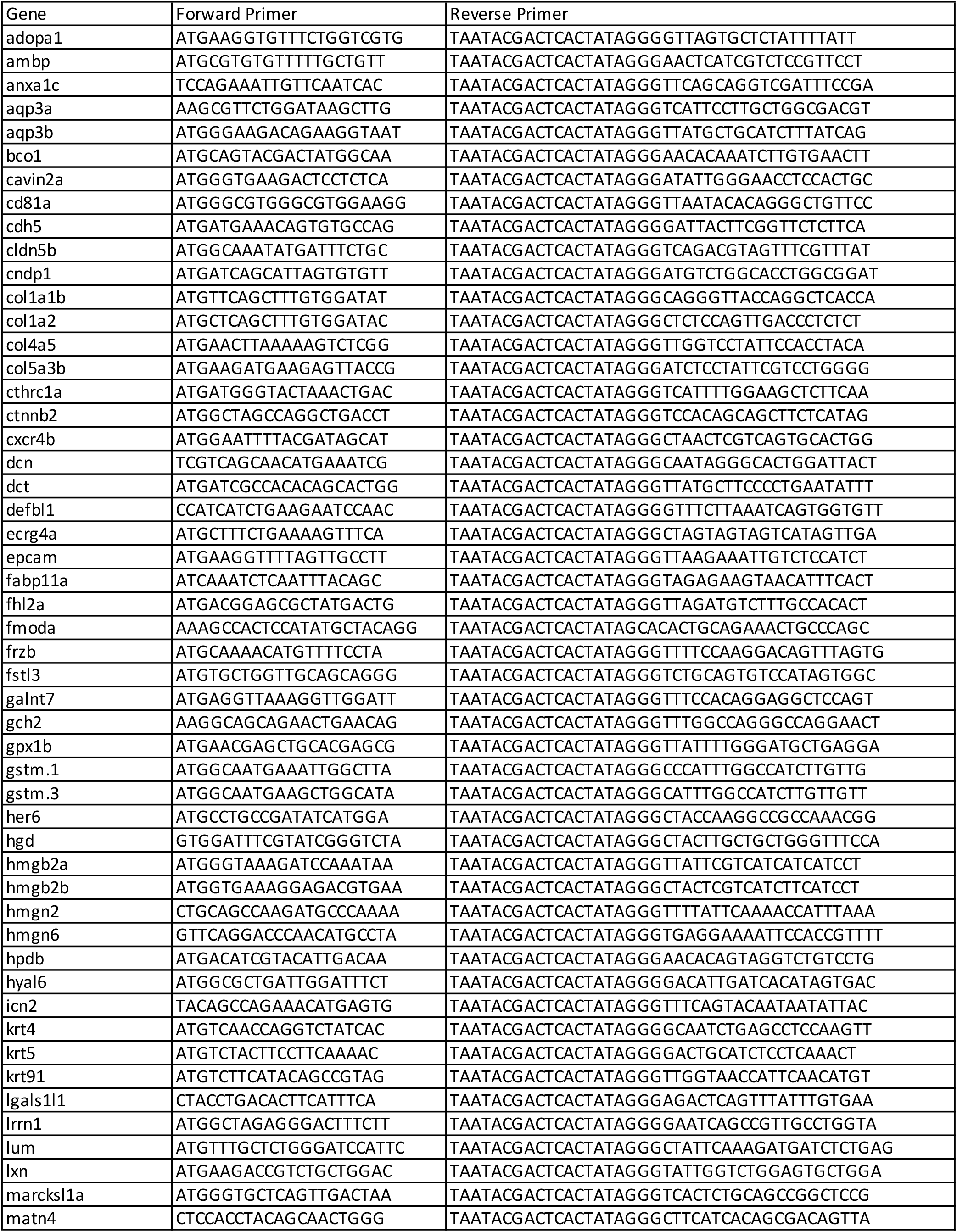
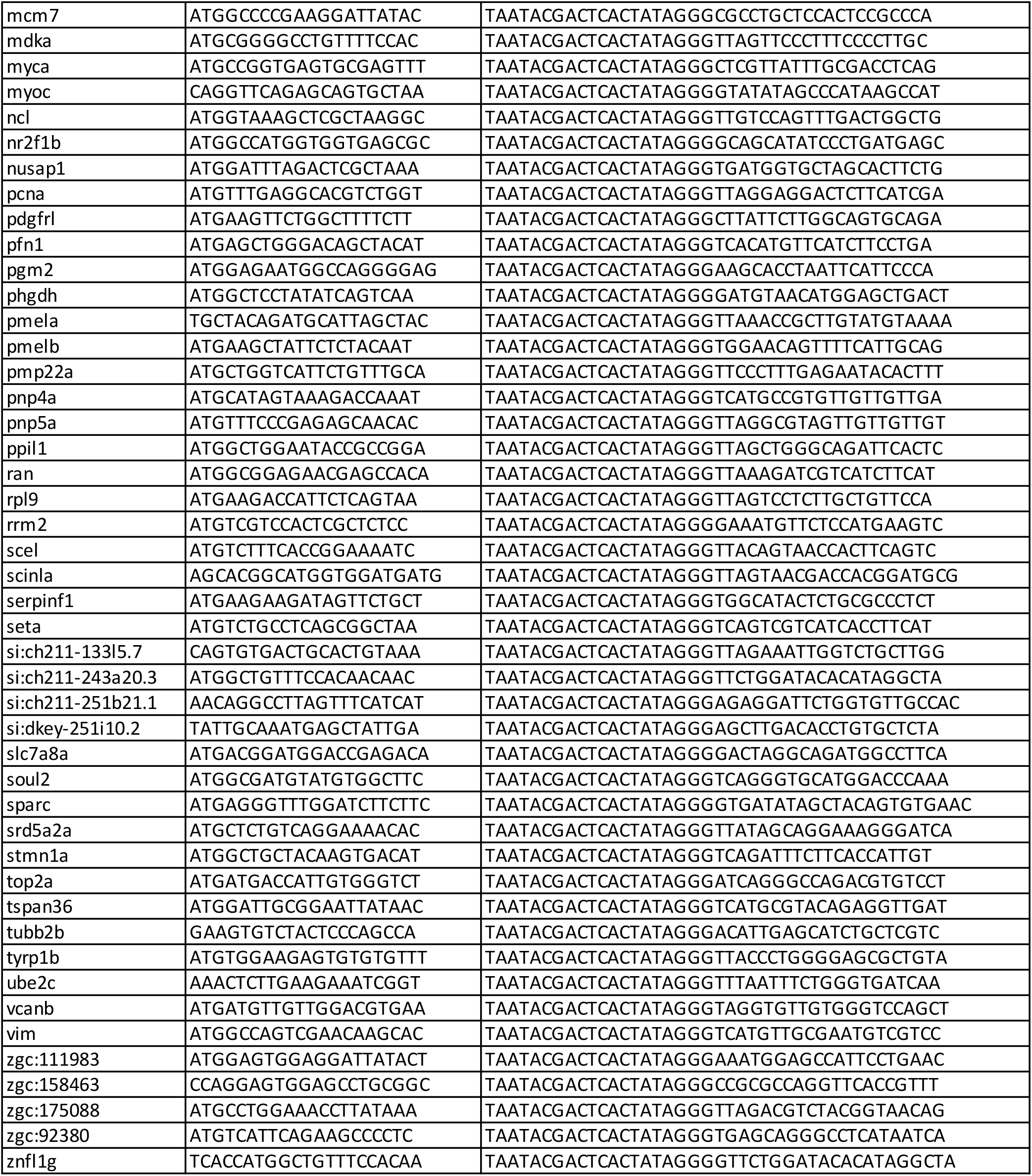

### Single Cell Transcriptome analysis

Single cell transcriptomes were generated as described previously (39). In brief, embryos from the transgenic line Tg[*foxc1b*:GFP] were collected every 24 hours between 48hpf and 144hpf. The embryos were anesthetized with 3-amino benzoic acid ethyl ester (Tricaine). Then, their eyes were dissected and subsequently collected on ice, before being incubated for 8 minutes in 0.25% Trypsin with EDTA at 37°C. After filtering, the dissociated single cells were sorted for GFP+ at the University of Kentucky Flow Cytometry and Immune Monitoring Core at the Markey Cancer Center. Thereafter, approximately 2000 cells per time point were loaded onto Chromium 10x V3 chips (10x genomics) and processed in the University of Kentucky Department of Biology Imaging Core to generate single cell barcoded cDNA. For sequencing, samples were sent to and processed at the University of Illinois at Urbana-Champaign Roy J. Carver Biotechnology Center. The resulting sequences were aggregated (incorporating mapped normalization), using the Cell Ranger 6.0 software (10x genomics). Further, we used the cell ranger pipeline to align our data to the GRCz11 assembly (NCBI Genome accession number: GCA_000002035.4).

Monocle 3 software was used according to the recommended settings to identify cell clusters, construct single-cell trajectories and identify differentially expressed genes (40). In brief, for every timepoint analyzed, cell types were clustered using PCA (n = 100 dimensions), we reduced the dimensions using UMAP and removed batch effects according to Haghverdi et al. (41). After clustering, we identified the top 50 over-represented genes per cluster and manually annotated the represented cell types, based on the findings from our *in situ* hybridizations and public databases. To perform pseudotime analysis, we aggregated our five timepoints, performed preprocessing clustering as described above for individual timepoints, and then manually identified roots using the convention that cell types from younger individuals should represent earlier stages in their developmental trajectory. Top 500 genes from each cluster at each time point are found in table 2.

### Fluorescent two color in situ hybridization (FWISH)

2 color fluorescent *in situ* wholemount hybridization (FWISH) was performed as previously described (39). In short, FITC and DIG labelled probes were co-hybridized, detected individually using FastBlue or FastRed (Sigma), and subsequently imaged using the C2+ Nikon confocal microscope using a 20X 0.95NA objective. DAPI was added to stain nuclei. 3D stacks were collected with 3μm steps and displayed as volume projections.

## RESULTS

### Assembly of a developmental AS transcriptomic atlas

To determine transcriptional changes occurring during early AS development we sought to isolate and analyze single cell transcriptomes of AS progenitors, the ASM. Owing to the lack of a universal genetic marker for ASM cells, analysis of whole eye tissue would limit our ability to isolate and specifically analyze ASM cells. As such, in order to facilitate direct sampling of ASM cells we employed the transgenic reporter line: Tg[*foxc1b*:GFP]. We have recently shown that ASM cells labelled in Tg[foxc1b:GFP] migrate into the AS space and contain transcriptomic profiles consistent with being AS progenitors (39). The *foxc1* transcription factor is also a well-known regulator and marker of the AS. To confirm that Tg[foxc1b:GFP] GFP+ cells can serve as reliable sources of ASM cell thorough AS development we assayed their expression patterns over AS development from 24-144hpf (Fig 1A). 3D confocal images from whole mount Tg[foxc1b:GFP] embryos show a robust and highly localized pattern of GFP+ cells in the anterior most regions of the eye, or the AS, up to and including 144hpf. Specifically, Tg[*foxc1b*:GFP] GFP+ cells were found throughout the central and peripheral regions of the AS at all the timepoints examined. 3D imaging confirmed that Tg[*foxc1b*:GFP] GFP+ cells populate the entire AS including the cornea and annular ligament. Overall, our previous and current characterizations indicate that Tg[*foxc1b*:GFP] GFP+ cells offer a comprehensive representation of ASM during the timepoints examined, 48-144hpf, and therefore provide an appropriate source for transcriptomic analysis of AS development. To generate ASM specific transcriptomes, we collected whole eyes of Tg[foxc1b:GFP] embryos at 48, 72, 96, 120 and 144hpf, dissociated them into a single cell suspension. Because Tg[*foxc1b*:GFP] GFP+ cells are not exclusive to the AS, FACS was performed on cells from dissected eyes, therefore ensuring only ASM associated GFP+ cells were collected. Purified ASM GFP+ cells were processed using the 10X genomics Chromium platform to generate scRNA libraries (Fig 1B). Two independent biological replicates were collected for each time point. Sequencing results indicated an average of ~2,400 cells total were captured per time point, with a total data set cell count of 12,234. Each of our samples had an average of 800+ genes identified from ~100,000+ reads/cell. Sequencing data was processed using CellRanger6.0 and subsequently analyzed for UMAP based cluster distribution and pseudotime using Monocle3.

**Figure 1:**
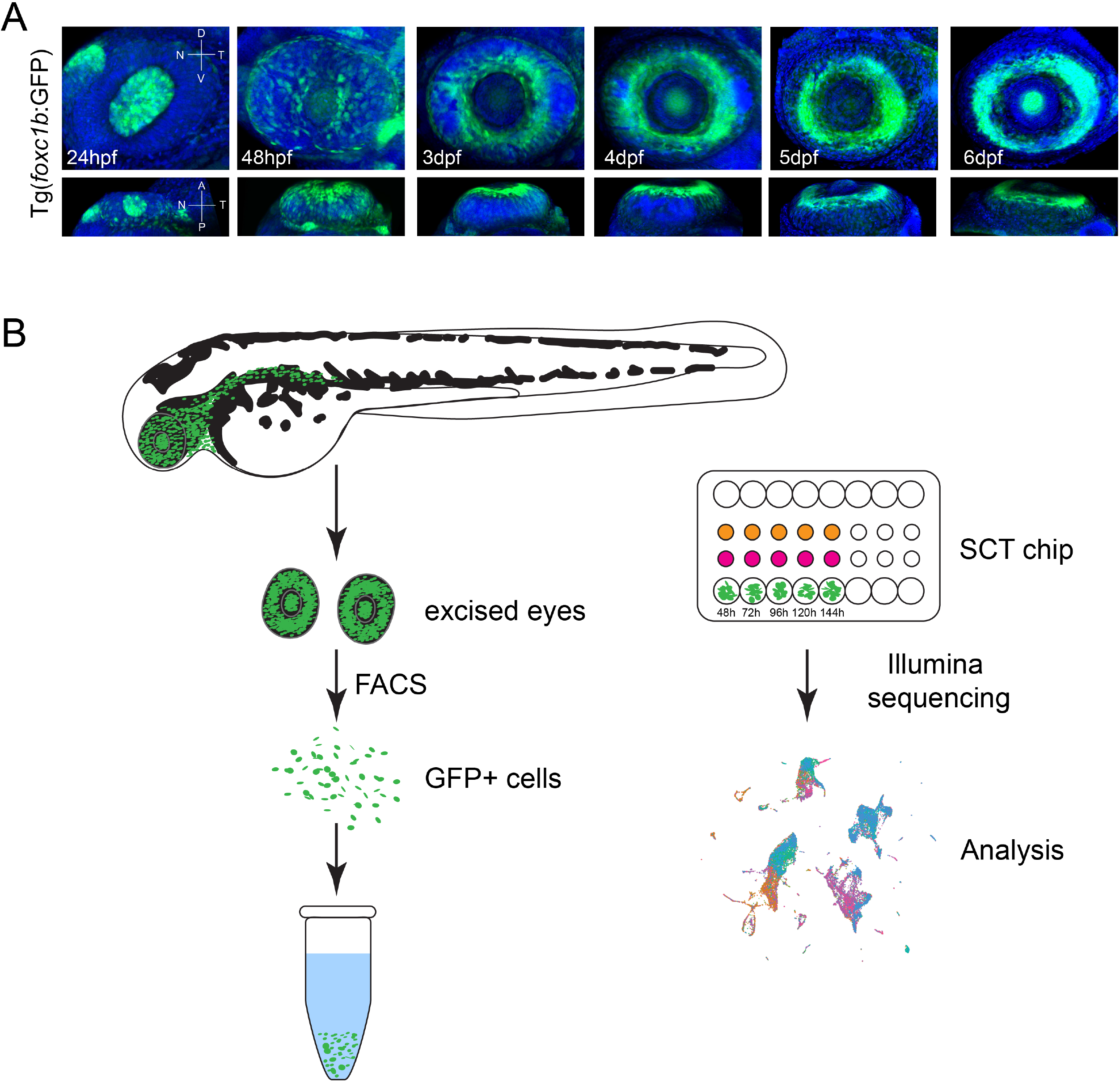
Strategy for generating ASM single cell transcriptomes over the course of early AS development. **A)** Volume projections of ventral and dorsal views from 3D confocal stacks of whole mount Tg[foxc1b:GFP] embryos at 24, 48, 72, 96, 120 and 144hpf. GFP+ cells (green) can be seen populating the AS regions by 48hpf and assembling the typical doughnut like AS localization pattern from 72-144hpf. DAPI (blue) was used to stain for nuclei. **B)** Schematic for capture and subsequent assembly of single cell ASM transcriptomes. Dissected whole eyes were dissociated and GFP+ cells isolated using FACS. Subsequently single cells are captured using the 10X chromium SCT chip and the resulting single cell cDNA sequenced and analyzed.

### Anterior segment tissue type classification

Prior to conducting cluster and pseudotime analysis of the ASM scRNA data, we sought to establish classification criteria for AS tissue types. Since whole eyes were dissociated to isolate GFP+ cells via FACS, we needed to establish criteria to identify which cell and tissue types are represented by the clusters. Although zebrafish are well established as a model for ocular development, the field lacks a comprehensive set of marker genes for developing AS tissues. To establish reliable genetic markers for corneal and AL tissues, we conducted extensive literature and databank (zfin.org) searches to classify genes for specific tissue types. Since our study is the first to characterize zebrafish developmental stages of AS formation, we relied on previous research that primarily focused on adult tissues from various species and extrapolated from those findings (3, 4, 18, 19, 28–32). As a primary source, we turned to recent single cell transcriptome analyses of individually dissected anterior segment tissues generated for various mammals, including human, different monkeys, and pigs (17–19, 29–32, 42). Additionally, we also referenced analyses of the zebrafish AS performed using 7dpf and adult tissues (43). Subsequently, we performed whole mount *in situ* hybridization (WISH) for chosen candidate genes at three different developmental stages, 24hpd, 3dpf and 5dpf, including cryosections to validate expression patterns (Fig 2). The chosen stages represent initiation of ASM colonization, completion of colonization and differentiation and finally rudimentary assembly of the AS respectively (9, 39). The two major tissue types belonging to the anterior segment we sought to classify were the cornea and the annular ligament (Fig 3). Both are known to derive from POM and therefore the ASM. The cornea we subdivided into corneal epithelium (CEp), corneal endothelium (CEn) and corneal stroma (CSt). We based classification of **CEp** on expression of keratins *krt4, krt5* and *krt91* as well as *icn2 (44–46)*. Among the studied tissues, the corneal epithelium is the most studied not only in zebrafish, but also mammals, likely due to its external position in the eye, which makes it easily accessible (3, 31, 32, 42). Also, as pointed out by Takamiya and co-workers (2015), penetrance of *in situ* probes can be more difficult in deeper tissues (43). For all the CEp marker genes, *krt4, 5, 91* and *icn2*, we can clearly distinguish expression across the anterior most corneal region, the epithelium (Fig 2A). Cryosections of 5dpf larvae further confirm these expression patterns (Fig 2B). Combinatorial expression analyses, using two color fluorescent WISH (FWISH) illustrates the high degree of co-expression of CEp markers at 5dpf (Fig 2C). We classified **CEn** identity based on the expression of *pmp22a* and *ctnnb2* (43, 47), which was confirmed via WISH and cryosections (Fig 2D-E). Expression of CEn markers differs from the CEp associated keratins with a significant reduction of expression in the most anterior corneal regions and higher expression in the dorsal and ventral periphery of the cornea. Maturation and differentiation of the corneal endothelium is a relatively slow process and full maturation is only reached after approximately two months (10). A recent publication described several marker genes of corneal tissues, however only two genes were described in the corneal endothelium *pitx2* and *ctnnb2* (43). Though, in the developing embryo the expression of *pitx2* is not limited to the endothelium (39) and the expression of *ctnnb2* was not known at stages younger than 7dpf. Our *in situ* hybridizations indicated, that *ctnnb2* is indeed expressed in the developing cornea and might serve as a marker for the developing corneal endothelium at early time points. Furthermore, fluorescent double *in situ* hybridizations show, that *ctnnb2* is co-expressed with *pmp22a* in the corneal endothelium of zebrafish (Fig 2F). Nonetheless, finding clear and specific marker genes for the corneal endothelium at these early developmental stages remains challenging due to ongoing differentiation. We classified **CSt** based on the expression of *dcn* and *lum* (11). The development of the corneal stroma is also a slow process and during the timepoints examined in this study must be considered mostly rudimentary and still maturing. Morphological studies of the stroma have shown how different collagens begin to form layers around 5-7 dpf, but they also describe the stroma itself as mostly absent of cells in these early stages (10, 20). WISH and cryosection images of *dcn* and *lum* confirm expression in the corneal region, primarily anterior of the lens (Fig 2G-H), while co-expression can be detected in ventral and dorsal regions (Fig 2I). As such, we determined these markers to be suitable for CSt identification (Fig 2G-H). Finally, we chose to classify the **AL** by the expression of the *genes hmgn2, hmgb2b, scinla*, as well as *myoc (18, 19, 48–51)*. As previously mentioned, the molecular composition of AL has been seldom studied in zebrafish and as such, apart from *scinla* (previously described as gsln1), much of our identification of markers of this structure was based on assumptions of conservation with the mammalian ICA. Previous studies in zebrafish mainly focused on identifying a specific marker but did not explore the gene expression profile in detail (49–51). Hence, due to the similarities in morphology and function, we used genes described in mammalian TM studies to identify basic markers. Hereby, we primarily relied on single cell transcriptome analyses that included isolated trabecular meshwork and Schlemm’s canal cells as well as TM cell culture studies (18, 19, 48, 52–55). As was done for all the corneal tissues, AL marker gene expression patterns were also confirmed with WISH and cryosections (Fig 2J-K). In general, we observed two distinct patterns of expression, one concentrated solely within the AL, *hmgb2b, hmgn2* and *myoc*, and the other exhibiting expression in the AL and in the CEn, *scinla* and *lxn* (homolog of rarres1). FWISH analysis indicated that all four marker genes are found to co-express in the AL to varying degrees. Taken together, we felt confident to use the aforementioned tissue identifier genes as the criteria for assigning CEp, CEn, CSt or AL identity to cell clusters in our ASM scRNA dataset.

**Figure 2:**
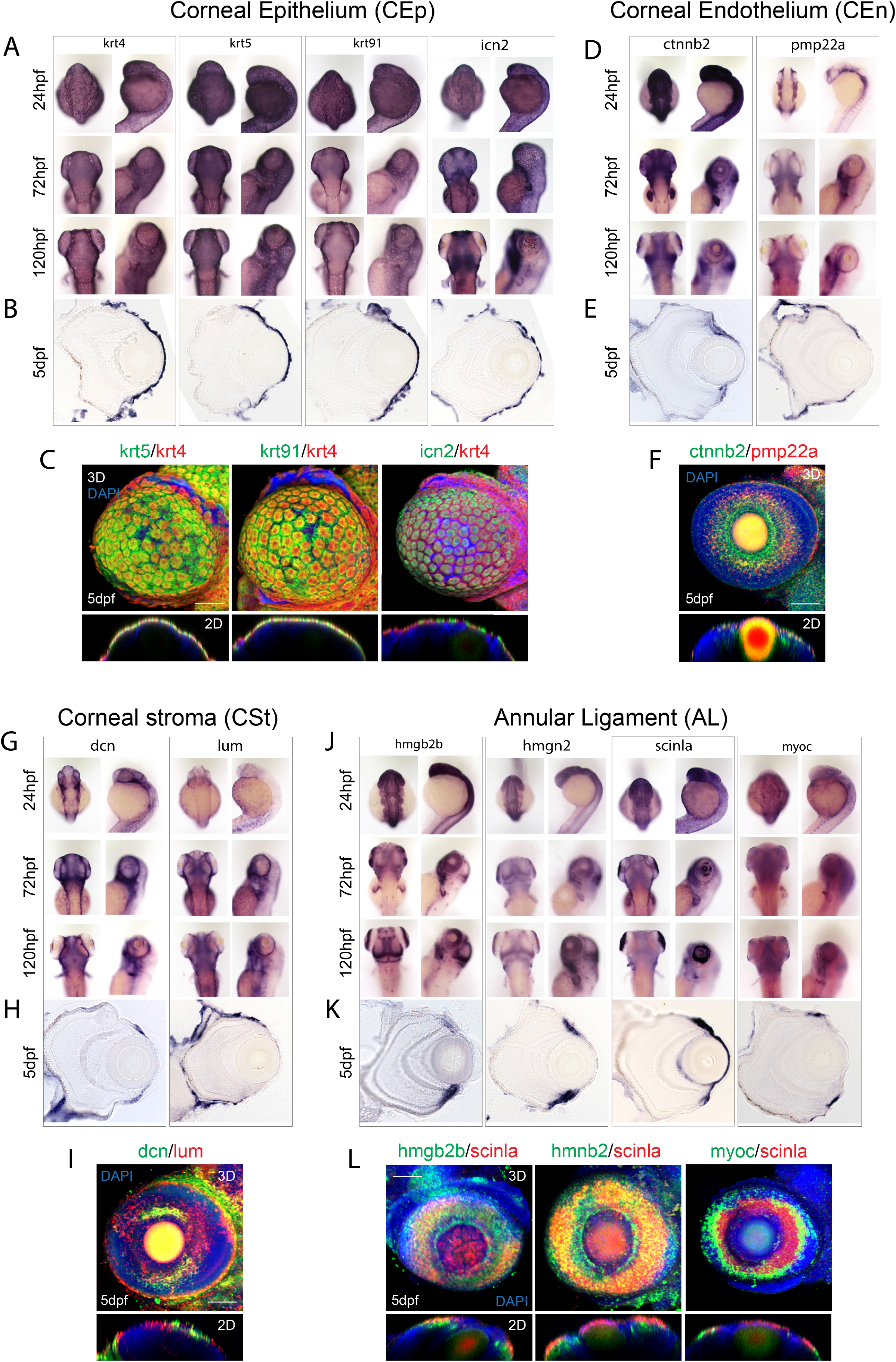
Assembly of markers for developing zebrafish corneal epithelium, endothelium, stroma and annular ligament. **A)** WISH analysis of corneal epithelium markers krt4, ktr5, krt91 and icn2 expression at 24, 72 and 120hpf displayed in dorsal and lateral views. **B)**10μm cryosections of 5dpf embryo eyes after WISH. **C)** Two color FWISH depicting co-expression of krt4(red)/krt5(green), krt91(green)/krt4(red) and icn2(green)/krt4(red) at 5dpf. DAPI (blue) was used to stain the nuclei. Volume projections in lateral view and single section dorsal views from 3D confocal stacks are displayed. Scale bar = 100μm. **D)** WISH analysis of corneal endothelium markers ctnnb2 and pmp22a expression at 24, 72 and 120hpf displayed in dorsal and lateral views. **E)**10μm cryosections of 5dpf embryo eyes after WISH. **F)** Two color FWISH depicting co-expression of pmp22a(red)/ctnnb2(green) at 5dpf. DAPI (blue) was used to stain the nuclei. Volume projections in lateral view and single section dorsal views from 3D confocal stacks are displayed. Scale bar = 100μm. **G)** WISH analysis of corneal stroma markers dcn and lum expression at 24, 72 and 120hpf displayed in dorsal and lateral views. **H)**10μm cryosections of 5dpf embryo eyes after WISH. **I)** Two color FWISH depicting co-expression of lum(red)/dcn(green) at 5dpf. DAPI (blue) was used to stain the nuclei. Volume projections in lateral view and single section dorsal views from 3D confocal stacks are displayed. Scale bar = 100μm. **J)** WISH analysis of annular ligament markers hmgb2b, hmgn2, scinla and myoc expression at 24, 72 and 120hpf displayed in dorsal and lateral views. **K)**10μm cryosections of 5dpf embryo eyes after WISH. **L)** Two color FWISH depicting co-expression of hmgb2b(green)/scinla(red), hmnb2(green)/scinla(red) and myoc(green)/scinla(red) at 5dpf. DAPI (blue) was used to stain the nuclei. Volume projections in lateral view and single section dorsal views from 3D confocal stacks are displayed. Scale bar = 100μm.

**Figure 3:**
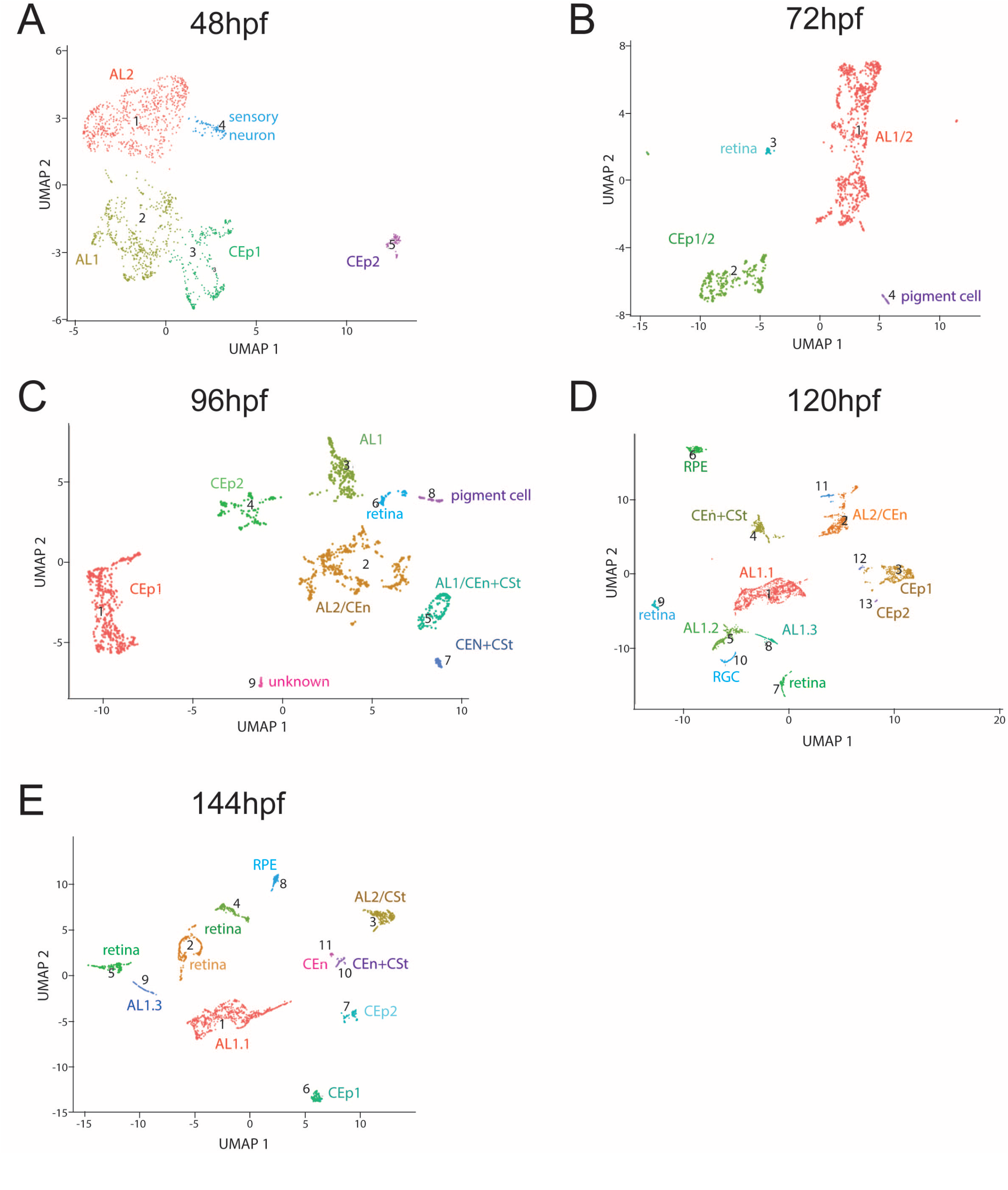
Clustering of purified ASM throughout early AS development. Cluster numbers are indicated in brackets. **A)** Cluster distribution at 48hpf. The five clusters observed were identified as: CEp1(2), CEp2(5), AL1(1), AL2(3) and sensory neurons(4). **B)** Cluster distribution at 72hpf. The four clusters observed were identified as: AL, CEp, retina and pigment cells. **C)** Cluster distribution at 96hpf. The nine clusters observed were identified as: CEp1(1), AL2/CEn(2), AL1(3), CEp2(4), AL1/CEn+CSt(5), retina(6), AL1/CEn+CSt(7), pigment cell(8), unknown(9). **D)** Cluster distribution at 120hpf. The fifteen clusters observed were identified as: AL1.1(1), AL2/CSt(2), CEp1(3), AL1/CEn+CSt(4), AL1.2(5), unknown(6), retina(7), AL1.3(8), retina(9), RGC(10), unknown(11). unknown(12), CEp2(13), unknown(14), unknown(15). **E)** Cluster distribution at 144hpf. The twelve clusters observed were identified as: AL1.1(1), retina(2), AL2/CEn(3), retina(4), retina(5), CEp1(6), CEp2(7), RPE(8), AL1.3(9), AL1/CEn+CSt(10), CEn(11), unknown(12).

### ASM scRNA cluster analysis throughout early AS development

Having collected single cell sequencing data from various timepoints we first sought to analyze cellular distribution at each timepoint individually. We aggregated the biological replicates and performed UMAP clustering using Monocle3. We analyzed every timepoint, using PCA (n = 100 dimensions) and removed batch effects according to Haghverdi et al. (41). Assignment of cluster identity was based on expression of the marker genes previously established (Fig 2). As expected, data from early timepoints, 48hpf and 72hpf, depicted only a handful of clusters (Fig 3A-B). At this early time of AS development, we expected the ASM to remain multipotent. However, already at 48hpf we can identify clusters with AL and CEp signatures (Fig 3A). The presence of CEp clusters was unexpected as corneal epithelium has been proposed to derive solely from the ectoderm (see discussion). Interestingly, CEp was bifurcated into CEp1, classified by elevated expression of *krt5* and *krt91*, and CEp2, classified by elevated expression of *krt4, cyt1* and *icn2* (Fig S1). The AL signature was also found in two distinct populations termed AL1 and AL2 where AL1 is designated by elevated expression of *hmgn2, hmgb2b, mcm7* and *pcna* while AL2 is designated by elevated expression of *scinla* and *lxn* (Figs 3A, S1). By 96hpf, we observed a significant stratification of clusters, in particular for those associated with the annular ligament (Fig 3C). Individual AL1 and AL2 clusters persist, but at 96hpf we additionally detected clusters with signatures of the CEn (elevated *pmp22a* and *ctnnb2* expression) mixed with AL2 (AL2/CEn) (Figs 3C, S3) and with CSt (CEn+CSt). This suggests that AL2, CEn and CSt cells may share a common progenitor and at 96hpf still retain some levels of multipotency. 96hpf is also when we detect the first clusters with signatures of solely CSt (elevated *dcn* and *lum* expression) mixed with CEn (Fig S3). Again, this suggests that cells slated to become AL, CSt and CEn share a common developmental pathway. Finally, we also continue to detect distinct CEp1 and CEp2 clusters at 96hpf. We never detect any association between CEp and AL, CEn or CSt clusters at any timepoints. As development proceeds, throughout 120 and 144hpf, we continued to detect all the aforementioned corneal and annular ligament clusters in addition to detecting the first sole CEn cluster at 144hpf (Fig 3D-E, S4-5). We never found a sole CSt cluster, likely stemming from the slow immigration of cells into these tissues which concludes later than 144hpf. To further validate the choice of genetic markers for assigning our cluster identities, we also examined relative expression of said markers at each timepoint (Fig S6). The majority of the assigned markers display a steady state expression throughout early development further confirming their suitability for tissue identification. Taken together, our data suggest that the early stratifications, between 72 and 96hpf, become established developmental pathways for these cell types. These conclusions are supported by morphological characterizations of the timing of AS development in zebrafish (8, 9). First cells within the iridocorneal angle can be identified as early as 3dpf, which grow in number at 5dpf. However, it was also described that a full histological identification of the annular ligament needed until 17dpf and hence, a full functional availability of these cells at early timepoints was questioned (9).

In addition to AS associated cell types, we also detected reoccurring clusters containing non-AS associated cells (Fig 3). This likely represents the fact that while the majority of Tg[*foxc1b*:GFP] GFP+ cells was isolated from whole eyes represent ASM cells, a small subset must also involve non-ASM cells. The most abundant non-ASM associated cells included cells of the cardiovascular system (BV), expressing the markers *ednraa, rasl12* and *tie1* (56, 57). They also included pigment cells (P), expressing genes such as *tyrosinase* and *gch2* (58, 59). The pigment cell clusters were found at 72 and 96hpf, and subsequently likely transition into retinal pigment epithelium (RPE) clusters throughout later timepoints (Fig 3). Pigment cells, as iris stroma, are known to include a NCC developmental pathway and therefore it is not surprising to find them in our data set. Expression of select genes verified to associate with pigment cell fates are summarized in supplemental figure 7. In addition to vasculature and pigment cells, we also identified retinal-like clusters which were found to contain different retinal cell types, including rods (marker: *rho*), cones (marker: *gnat2*) and retinal ganglion cells. In total we found 21 clusters in our combined dataset. However, it must also be stressed that most non-ASM associated clusters contained very few cells compared to those of corneal or annular ligament lineages.

### Molecular characterization of AS clusters during early development

To showcase the unique molecular differences associated with each of the identified AS lineages (AL1, AL2/CEn, CEn+CSt, CEp1 and CEp2) we examined the top upregulated genes from each cluster at each timepoint (Fig 4). At 48hpf, as expected, the AL associated clusters exhibit high levels of expression for *hmgb2b and hmgn2*, but also *hmgb2a, h3f3a, fabp7a, cfl1, ccnd1* and *tmsb4x* (Fig 4A). Uniquely for AL2 we also observe *pmp22a, tgfbi* and *tpm4a* upregulation. CEp1 and 2 are both defined by very high levels of *krt4* expression (Fig 4A). CEp1 also exhibits higher levels of collagens *5a3b* and 1a1b as well as *pnf1* compared to CEp2, while CEp2 exhibits high levels of *cyt1, cyt1l, anxa1c, agr1* and *icn2* (Fig 4A). At 72hpf, the primary difference between the AL and CEp multipotent clusters is high expression of krt4, epcam, cyt1 and tmsb1 in CEp and *hmgn2, cd81a, mdka, si:ch211-133n4.4* and *pmp22a* in AL (Fig 4B). At 96hpf, the time of significant stratification of the clusters, we see a much more defined molecular signature for all the different tissue types. CEp1 is defined by high expression of *krt4, krt5, krt91, pfn1, cldni* and *col11a1a* while CEp2 exhibits unique upregulation of *malb, cldnb, cldne, krt1-c5, cldnh* and *tnks1bp1* (Fig 4C). AL1 is defined by high expression of *mdka, ccnd1, cadm3, hmgn2, fabp7a, mcm7, cad* and *hmgb2b*. CEn+CSt on the other hand exhibits high *dcn, col5a2a, cvanb, sema3d, scara5, osr1, add3a, pmp22a* and *cd81a*, while AL2/CEn cells display upregulated expression of *myl9a, fstl3, pitx3, hgd, foxc1a, pitx2* and *fmoda* (Fig 4C). At 120hpf, CEp1 continues to express *krt4, krt5, krt91, pfn1, cldni* at high levels, while also increasing expression of *krt8*. CEp2 also exhibits high expression of *krt4, krt5, krt91*, and *pnf1*, in addition to *icn2*, anxa1c and several unknown genes including *zgc:193505, si:ch1073-340i21.3* and *si:ch211-157c3.4* (Fig 4D). By 120hpf AL1, comprised of 3 individual clusters, AL1.1, AL1.2 and AL1.3. All three are defined by continued high expression of *hmgn2, hmgn6* and *hmgb2a* while upregulation of *nova2, gpm6aa* and *ccnd1* defines AL1.2 and AL1.3. CEn+CSt displays upregulation of *rn7sk, pmp22a, cd81a* and *krt8*. Distinct upregulated expression in AL2/CEn includes *pitx2, pitx3, scinla, fmoda* and *fxyd1* (Fig 4D). By 144hpf, we continue to see similar top upregulated genes in CEp1 while CEp2 exhibits upregulation of *agr1, tcnbb, icn2* and unknown genes *si:ch211-195b11.3* and *zgc:193505* (Fig 4E). AL1.1 and AL1.2 show similar expression patterns with upregulated *hmgb2a, mcm7*, but also now upregulate *cadm3, anp32e* and *mdka*. CEn+CSt continues to express high levels of *pmp22a*, while also increasing expression of *matn4, col9a2, col2a1a, cnmd* and *irx1b*. AL2/CEn also continues to express *pmp22a* at high levels, in addition to *hpdb, pitx3, hgd* and *scinla* (Fig 4E). The first independent CEn cluster is defined by upregulated expression of *marcks1a, pmp22a, scl43a3b, cyp1ad3* and *foxc1b* (Fig 4E). Based on these expression patterns we observe that many of the genes used in our assay as determinants of cell fate are constantly expressed at high levels throughout early development while other genes, many previously not associated with AS development, display various patterns of expression.

**Figure 4:**
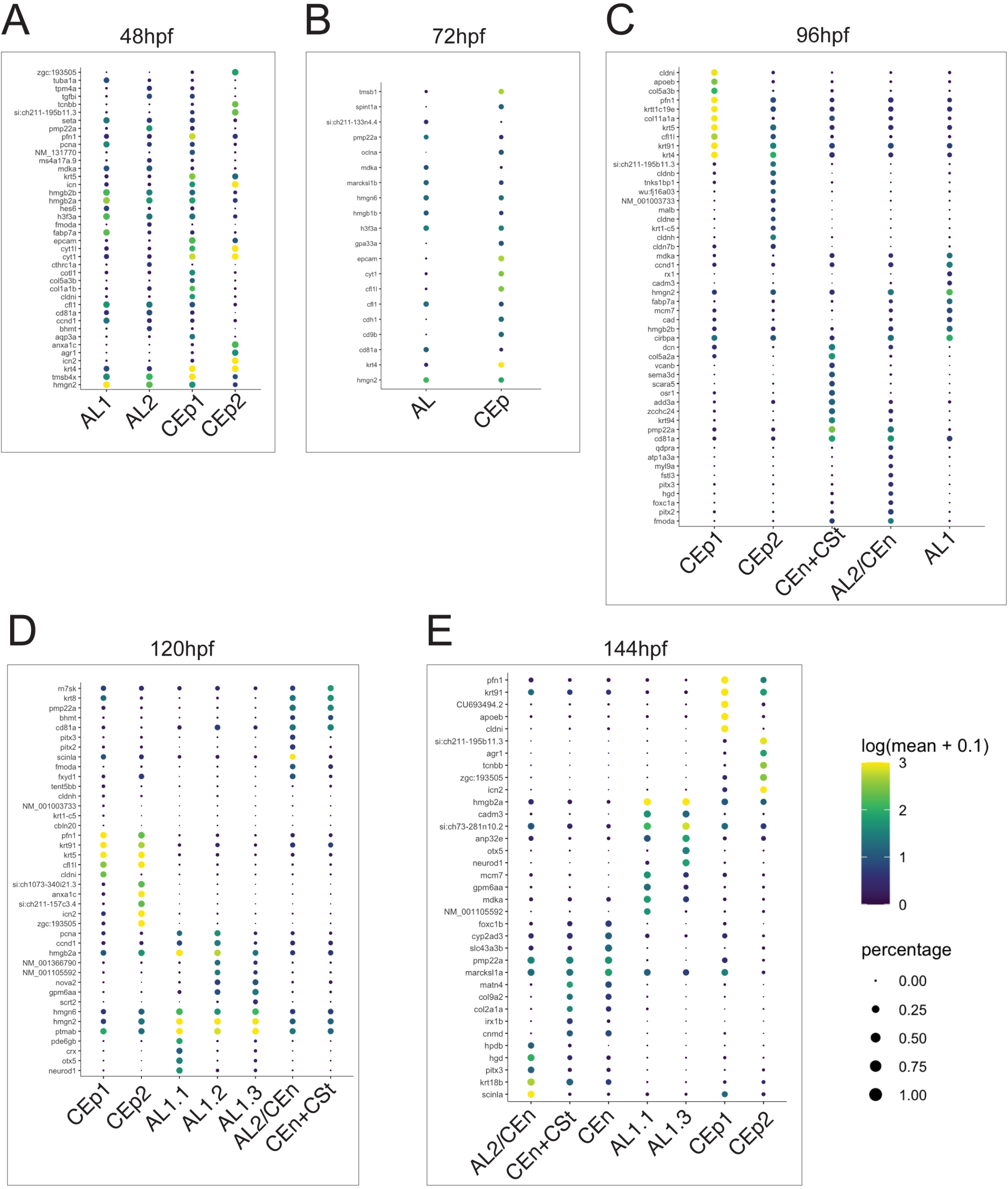
Top 10 genes for AS related cell types at each timepoint examined. **A)** Top10 genes from AL1, AL2, CEp1 and CEp2 at 48hpf. **B)** Top10 genes from AL and CEp clusters at 72hpf. **C)** Top10 genes from AL1, AL1/CEn+Cst, AL2/CEn, CEp1 and CEp2 clusters at 96hpf. **D)** Top10 genes from AL1.1, AL1.2, AL1.3, AL1/CEn+CSt, AL2/CEn, CEp1 and CEp2 at 120hpf. **E)** Top10 genes from AL1.2, AL1.3, AL1/CEn+CSt, AL2/CEn, CEn, CEp1 and CEp2 at 144hpf. Scale of degree of expression and percentage is indicated.

### Developmental dynamics and trajectory analysis of AS development

Having classified AS cell types over the course of early AS development, we next sought to track trajectories and relationships in time. To establish and illustrate the developmental relationship between clusters over time, we compiled a timeline (Fig 5). Trajectories were determined by manually examining the top50 upregulated genes in each cluster and comparing patterns of expression in subsequent time points to generate most likely trajectories over time. Starting at 48hpf with the AL clusters, AL1 and AL2, we track their convergence into one multipotent cluster at 72hpf. By 96hpf we track the AL cell type to stratify into AL2/CEn, AL1 and CEn+CSt. The main AL1 lineage further stratifies into three sub clusters at 120hpf, AL1.1, 1.2 and 1.3, and settles as two clusters, AL1.1 and AL1.3 by 144hpf. The CEn+CSt lineage also appears to be specified by 96hpf and persists up to 144hpf. However, at 144hpf we also see a portion of CEn+CSt diverge into a possibly committed CEn lineage. These observations indicate that the AL1 lineage exhibits multipotency including potential for CEn and CSt lineages early in development. Conversely, starting at 96hpf, the AL2/CEn cluster persists up to 144hpf and is therefore likely already specified by 96hpf. This suggests a very close relationship between AL and CEn cells during early development. When tracking CEp development, starting with CEp1 and CEp2, we observe the two clusters converge at 72hpf. However, this convergence may be the result of generally fewer CEp2 cells which may be masked by the large number of CEp1 cells at this timepoint. Subsequently, they re-diverge into CEp1 and CEp2 by 96hpf and both trajectories remain independent up to 144hpf. Overall, during early AS development the number of clusters representing the corneal epithelium remains at two, indicating the slow developmental pace after their rapid initiation. This finding is supported by the fact that zebrafish corneal epithelium remains 2-layered for the first four weeks (10).

**Figure 5:**
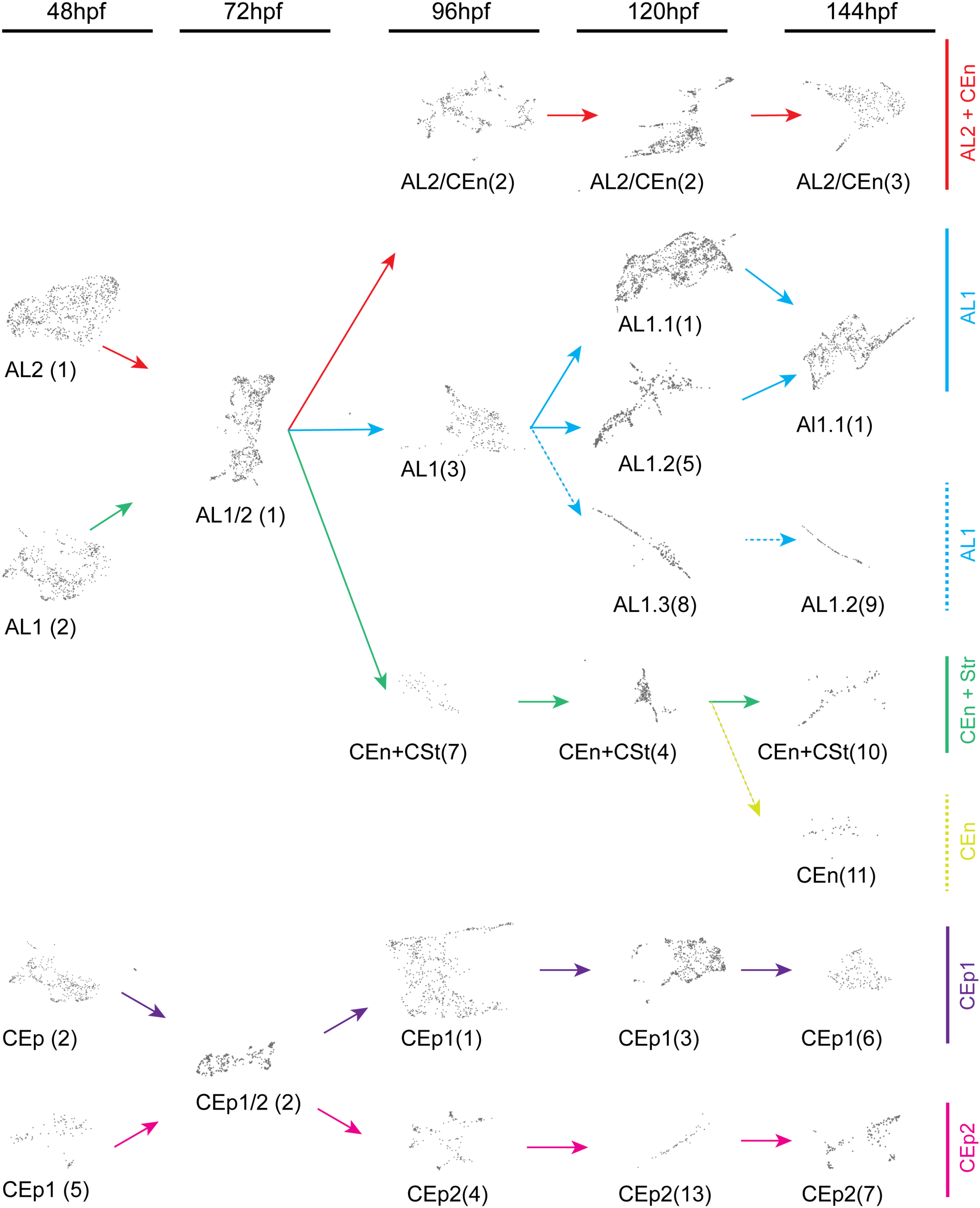
Cluster trajectories during early development of corneal epithelium, endothelium stroma and annular ligament. Developmental trajectories of individual clusters from each timepoint, 48-144hpf, displaying temporal relationships. Numbers in brackets correspond to timepoint specific clusters (Fig 3). Arrows indicate lineage relationships of the AL1 (blue), AL2/CEn (magenta), AL1/CEn+CSt (blue), CEn (light blue), CEp1 (purple) and CEp2 (pink) cell types. CEp lineages show early and dedicated commitment while AL lineages display multipotency for AL, CEn or CSt cell types.

To complement our cluster-based trajectories over developmental time we also performed a pseudotime analysis of our combined dataset using Monocle3. The major difference from our cluster analysis is that pseudotime analysis focuses on the trajectory of cells instead of whole clusters and helps identify how gene expression changes within a cell trajectory. Hence, this analysis allowed us to characterize if particular marker genes are strongly expressed at specific points in the trajectory. To perform pseudotime analysis all our timepoints were aggregated into a single data set and then subsequently analyzed using the Monocle3 pseudotime algorithm (Fig 6A, S8). We summarize our results based on the AS tissue types being tracked.

**Figure 6:**
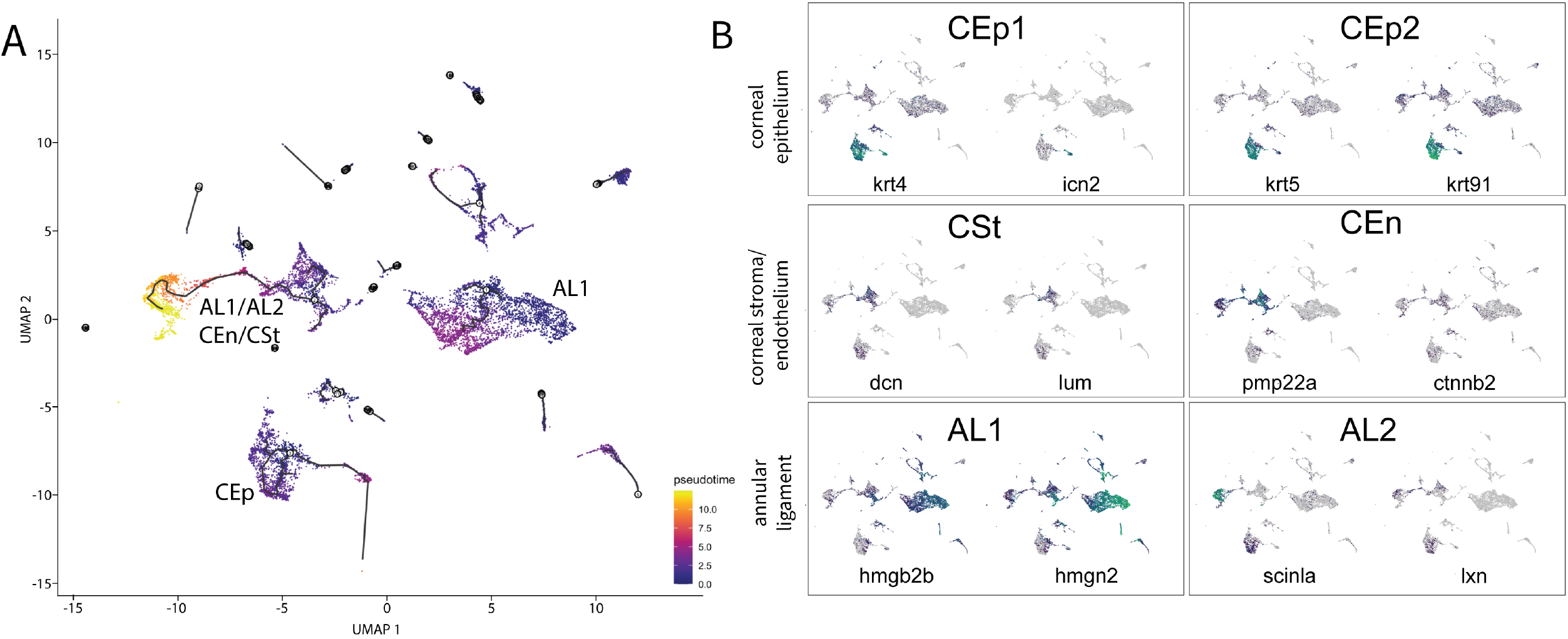
Pseudotime analysis of purified ASM cells during early AS development. **A)** The trajectory analysis showing three major trajectories including CEp, AL+CEn/CSt, AL1. **B)** Marker genes for the different cornea tissues and annular ligament projected onto the trajectory graph. Markers for CEp1: krt4 and icn2, for CEp2: krt5, krt91, for CSt: dcn and lum, for CEn pmp22a and ctnnb2, for AL1: hmgb2b and hmgn2 and for Al2: scinla and lxn.

#### Corneal epithelium

When tracking markers of the corneal epithelium it becomes apparent, that marker genes representing CEp1 and CEp2 are expressed in this trajectory, indicating a close relation between the clusters. Many, of the characterized marker genes such as *krt4, krt5, krt91 or icn2* are strongly expressed throughout the identified trajectory, validating their general importance for the cells (Fig 6B). This appears to confirm the relatively conservative use of this core group of genes as markers for this tissue throughout early AS development. Other CEp associated genes display varying pattens of expression throughout the trajectory. For instance, *sparc* and *lgals1l1*, are strongly upregulated in the early parts of the trajectory but mostly absent from the later points (Fig S8A). Interestingly, in contrast to most keratins that are upregulated throughout the trajectory, collagens (*col1a1b, col5a3b, col17a1a*) are upregulated early, but then decrease over time (Fig S8B). Other genes, such as *icn2, anxa1c* and *pnp5a* show a reduced or completely absent expression in the early time points but are highly upregulated at the later points of the trajectory (Fig S8C).

#### Corneal endothelium and stroma

Genes marking the corneal endothelium were found in the same trajectory as the AL markers, confirming our cluster analysis results (Fig 6B). By analyzing the distribution of the genes in the trajectory, we found that *pmp22a* is strongly expressed throughout the entire trajectory. *Ctnnb2* and *dcn* on the other hand, share a sparse expression and a distribution that is relatively even throughout, but severely reduced toward the tail end of the trajectory (Fig 6B).

#### Annular ligament

When comparing the pseudotime distributions of AL marker genes, we found that both tissue types, AL1 and AL2 share the same trajectory, further confirming the close relation between the two (Fig 6B). AL marker genes *hmgb2b* and *hmgn2* are broadly distributed throughout the AL trajectory. However, upon closer inspection it appears that *hmgb2b* is present in early sections of the trajectory while mostly absent from later branches (Fig 6B). *Scinla* and *fabp11a* (also called *fabp4a*) on the other hand, are mostly absent from the early branches of the trajectory but are strongly upregulated in later branches (Fig 6B, S8D). Two previously unknown AS associated genes, *hgd* and *si:ch211-251b21.1* as well as *vim* and *hpdb* can be found expressed early in the trajectory and then again towards the end of the trajectory (Fig S8E), while *fmoda* can be found throughout all branches of the trajectory (Fig S8F). As such, our analysis of AL development, the first to be conducted in zebrafish, clearly defines this tissue as having arisen from multipotent progenitors which over time specify in AL, CEn and CSt dedicated populations. Overall, pseudotime analysis supports our cluster-based trajectory tracking and establishes developmental trajectories for zebrafish AS tissues.

### Conservation of early zebrafish AS development

Considering our data set being the first to examine expression over the course of AS development we wanted to assess the utility of our data set. To do so, we referenced genes in our data set to those that have been identified in mammalian, and particularly human AS tissues. Recent work examining adult cornea and the entire AS, also at single cell levels, was used to cross reference with our own (16–19, 29–32, 47). In doing so, we identified significant overlap in conservation of expression for CEp, CEn as well as the AL, compared to the cornea and trabecular meshwork, respectively. Using WISH, cryosections and two color FWISH we confirmed CEp expression patterns for 14 genes found within the CEp1 and CEp2 clusters (Figure 7A-B, S9A) that were also identified in human corneal data sets. We show clear co-expression of *zgc92380, sparc, pfn1, epcam* and *col1a1b* with our established CEp marker *krt4* (Fig 7C). This outcome thus suggests a high degree of conservation between zebrafish and human corneal epithelial development. In addition, we characterized expression of eight genes previously not associated with CEp development including: *cavin2a, col4a5, col5a3b, pnp5a, soul2, si211:133157, zgc:158463* and *zgc:175088* (Figure S9B). All of which display characteristic expression patterns for corneal epithelium. Furthermore, thanks to the multi timepoint nature of our data, we also analyzed total expression changes for each of these genes during the course of development by comparing relative expression within clusters across developmental time (Fig 7D). When examining *zgc92380, sparc, pfn1, epcam* and *col1a1b* relative expression levels in CEp1 we see a relatively steady state of expression throughout development with only *epcam* displaying a significant drop in expression at 96hpf (Fig 7D). In CEp2 we see a significant spike in expression of all five genes at 72hpf and a subsequent decline. For CEn associated genes, we confirmed the expression of *matn4*, a known component of the human cornea, and identified *cthrc1a* as a potential novel regulator of CEn development and or function (Fig 7E-F) (60, 61). Both display spatial expression patterns like those of CEn markers *pmp22a* and *ctnnb2*. When tracking relative expression levels over time, we note that in case of the CEn markers, pmp22a is initially relatively strongly expressed, but its expression decreases a 120hpf and slightly rebounds at 144hpf. Whereas the expression of ctnnb2 is relatively low and further decreases at 96hpf (Fig 7G). Matn4 expression declines by 96hpf with a slight rebound by 144hpf. Cthrc1a follows a similar pattern with a gradual decline, most evident at 96hpf, and a rebound by 144hpf (Fig 7G).

Very few genes were previously known to be expressed in the AL of zebrafish (49–51). Of those known, such as *scinla*, these genes were not specific markers of the annular ligament, as they are also found expressed in adjacent tissues, such as the corneal endothelium. Hence, this study provides the first extensive gene expression analysis of the zebrafish annular ligament and establishes a set of marker genes more specific to this tissue. Using WISH, we confirmed expression of 31 genes found in AL1 and AL2 clusters that are also known to be expressed in human TM (Figure 8 + S10A). Of those 31, ten were further characterized using cryosection as well as co-expression analysis with *scinla* using two color FWISH. Five of the genes characterized were associated with AL1, which included *nusap1, frzb, cxcr4b, rrm2* and *cdh5* (Fig 8A-B), while the other five, *mcm7, vim, lxn, fmoda* and *fabp11a* were associated with AL2 (Fig 8E-F). Cryosection images confirm the distinction as AL2 genes tend to display both AL and corneal endothelium expression patterns (Fig 8B, F). Furthermore, 3D and 2D confocal images clearly confirm co-expression of these genes with *scinla* in AL tissues (Fig 8C, G). When tracking mean expression within AL clusters over time, we observe steady state levels of expression for our canonical markers *hmgn2, hmgb2b, scinla* and *myoc* with a slight increase at 72hpf and 144hpf (Fig 8D). In case of *rrm2*, the expression increases constantly, but slightly drops at 144hpf. *Nusap1* on the other hand, is initially relatively weakly expressed, but strongly increases after 96hpf (Fig 8D). The expression of both, *mcm7* and *fabp11a* increases until 96hpf, but while mcm7 initially drops at 120hpf and then rises again, *fabp11a* remains at a constant level (Fig 8H). *Fmoda* and *rn7sk* are both expressed up to 72hpf, but their expression decreases in the following time points (Fig 8H). In addition to AL genes conserved in TM tissues, we also identified 13 potentially novel components of the AL (Fig 9+S10B). These included *stmn1a, si:ch211-251b21.1, hgd, phghd* and *cndp1* (Fig 9A). All five of these genes display AL1 characteristic expression, strong staining in the AL regions and absence from the cornea and co-expressing with *scinla* as shown using FWISH (Fig 9A-C). When tracking their mean expression patterns over time we note that *cndp1, phgdh* and *hgd* display steady state expression, *stmn1a* expression gradually increases over time while *si211-251b21.1* peaks at 72hpf and gradually declines thereafter (Fig 9D). Taken together, we not only confirm significant conservation of gene expression between human TM and developing zebrafish AL, but we also identify several new target genes as potential constituents of the AL. While we examined a large number of genes from our data set, many more remain to be validated using WISH and additional techniques.

**Figure 7:**
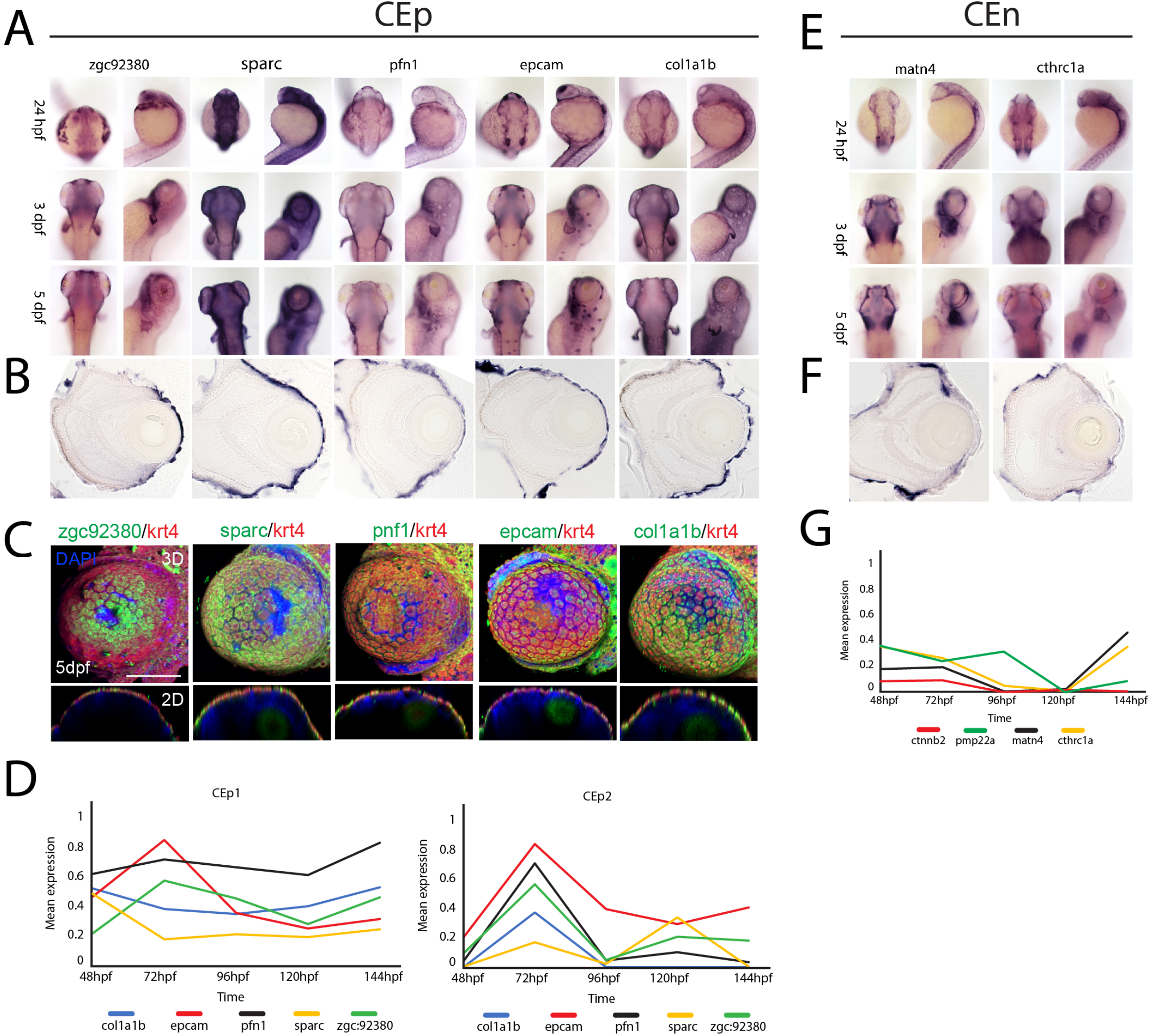
Spatiotemporal expression patterns of conserved corneal epithelium and endothelium associated genes. **A)** WISH analysis of corneal epithelium associated genes zgc:92380, sparc, pfn1, epcam and col1a1b expression at 24, 72 and 120hpf displayed in dorsal and lateral views. **B)**10μm cryosections of 5dpf embryo eyes after WISH. **C)** Two color FWISH depicting co-expression of krt4(red)/zgc:92380(green), sparc(green)/krt4(red), pfn1(green)/krt4(red), epcam(green)/krt4(red) and col1a1b(green)/krt4(red) at 5dpf. DAPI (blue) was used to stain the nuclei. Volume projections in lateral view and single section dorsal views from 3D confocal stacks are displayed. Scale bar = 100μm. **D)** Average mean expression measurements from CEp1 and CEp2 clusters for marker genes krt4, krt5, krt91 and icn2 as well as col1a1b, epcam, pfn1, sparc and zgc92380 at each timepoint examined, 48-144hpf. **E)** WISH analysis of corneal endothelium associated genes matn4 and cthrc1a expression at 24, 72 and 120hpf displayed in dorsal and lateral views. **F)**10μm cryosections of 5dpf embryo eyes after WISH. **G)** Average mean expression measurements from CEn clusters for pmp22a, ctnnb2, matn4 and cthrc1a at each timepoint examined, 48-144hpf.

**Figure 8:**
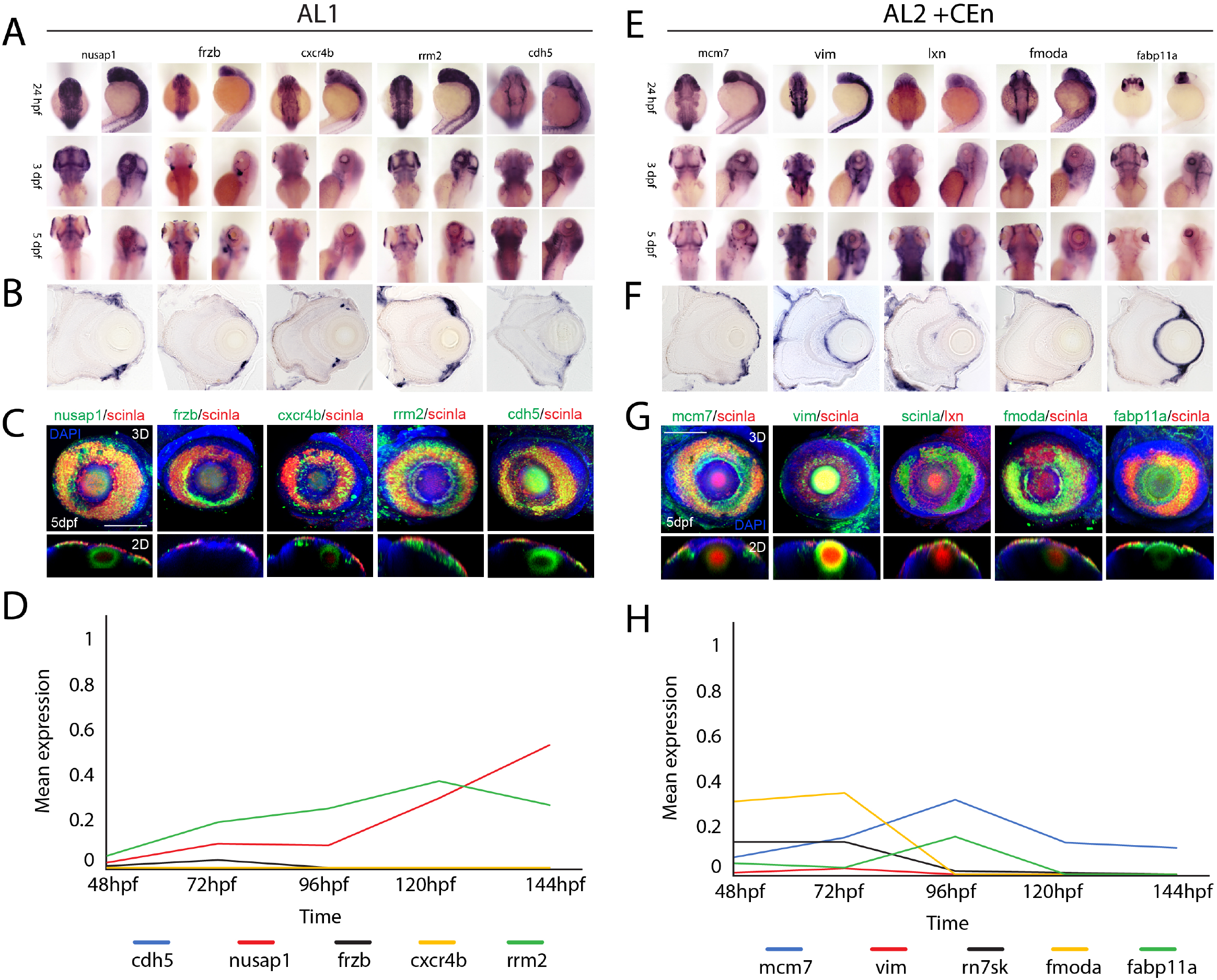
Spatiotemporal expression patterns of conserved annular ligament associated genes. **A)** WISH expression analysis of annular ligament 1 (AL1) associated genes *nusap1, frzb, cxcr4b, rrn2* and *cdh5* at 24, 72 and 120hpf displayed in dorsal and lateral views. **B)**10μm cryosections of 5dpf embryo eyes after WISH. **C)** Two color FWISH depicting co-expression of nusap1(green)/scinla(red), frzb(green)/scinla(red), cxcr4b(green)/scinla, rrn2(green)/scinla(red) and cdh5(green)/scinla(red) at 5dpf. DAPI (blue) was used to stain the nuclei. Volume projections in lateral view and single section dorsal views from 3D confocal stacks are displayed. **D)** Average mean expression measurements from AL1 clusters for genes *nusap1, frzb, cxcr4b rrn2*, and *cdh5* at each timepoint examined, 48-144hpf. **E)** WISH expression analysis of annular ligament 2 (AL2) associated genes *mcm7, vim, lxn, fmoda* and *fabp11a* at 24, 72 and 120hpf displayed in dorsal and lateral views. **F)** 10μm cryosections of 5dpf embryo eyes after WISH. **G)** Two color FWISH depicting co-expression of mcm7(green)/scinla(red), vim(green)/scinla(red), lxn(green)/scinla, fmoda(green)/scinla(red) and fabp11a(green)/scinla(red) at 5dpf. DAPI (blue) was used to stain the nuclei. Volume projections in lateral view and single section dorsal views from 3D confocal stacks are displayed. Scale bar = 100μm. **H)** Average mean expression measurements from AL2 clusters for genes *mcm7, vim, lxn, fmoda* and *fabp11a* over developmental time.

**Figure 9:**
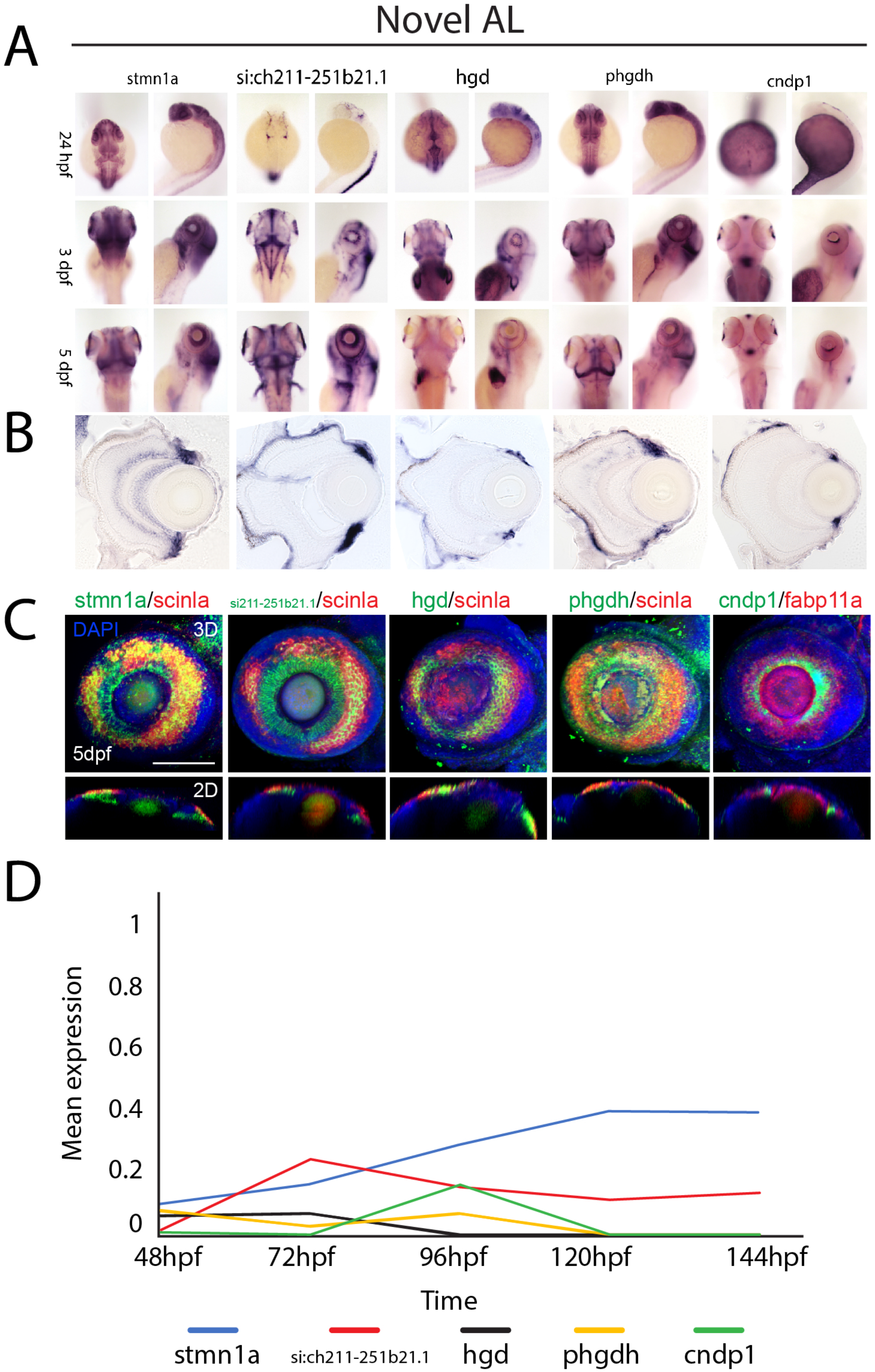
Expression patterns of novel annular ligament associated genes. **A)** WISH expression analysis of annular ligament associated genes *stmn1a, si:211-251b21.1, hgd, phgdh* and *cndp1* at 24, 72 and 120hpf displayed in dorsal and lateral views. **B)**10μm cryosections of 5dpf embryo eyes after WISH. **C)** Two color FWISH depicting co-expression of stmn1a(green)/scinla(red), si:ch211-252b21.1(green)/scinla(red), hgd(green)/scinla, phgdh(green)/scinla(red) and cndp1(green)/scinla(red) at 5dpf. DAPI (blue) was used to stain the nuclei. Volume projections in lateral view and single section dorsal views from 3D confocal stacks are displayed. Scale bar = 100μm. **D)** Average mean expression measurements over developmental time.

## DISCUSSION

In this study, we present a spatiotemporal analysis of 12,234 single cell transcriptomes of anterior segment tissues of zebrafish isolated during initial AS formation. While recent single cell transcriptomic studies focused on the analysis of mature tissues, our analysis focuses on early development. This gives us the unique opportunity to characterize and follow gene expression changes throughout multiple timepoints of early AS development. The key points of our analysis are the establishment of a defined set of marker genes for developing anterior segment tissues in zebrafish, the identification of multiple cell types in the cornea and annular ligament of zebrafish, establishment of developmental trajectories for corneal and annular ligament lineages and validating conservation of zebrafish AS gene expression with mammalian and human AS tissues.

### Establishing a set of marker genes for anterior segment structures in zebrafish

The focus of our analysis were the cornea and the annular ligament. Since developmentally the molecular composition of both of these tissues in zebrafish, especially the annular ligament, was limited to a handful of associated genes, we first had to establish a solid base of tissue marker genes. For the corneal epithelium we found that the keratins *krt4,5, 91* and also *icn2* to be excellent markers, which for the most part have also described in adult tissue of several other species (44–47, 62, 63). For the corneal endothelium we found that *ctnnb2* as well as *pmp22a* are already expressed in the early corneal tissues, confirming the finding of *ctnnb2* in zebrafish at later stages (43, 47). Additionally, *lum* and *dcn* are expressed in the developing corneal stroma, as reported in other organisms (11). Regarding the annular ligament, we identified *hmgb2b, hmgn2, mcm7, pcna* as well as *scinla* and *lxn* to be suitable expressed markers, of which only *scinla* was previously known to be expressed in the zebrafish tissue (49–51). Establishment of genetic markers for developing zebrafish AS tissues not only helps with their identification in transcriptomic data sets, but also enables comparisons to other model organisms. It also serves as a resource for performing subsequent unbiased analysis on cells isolated from whole eye tissues.

### Identification of Different cell types in the cornea and annular ligament

Analysis of our combined dataset revealed a total of 21 clusters, indicating the presence of multiple cell types. AS related cell types included the annular ligament, corneal endothelium, corneal stroma and also corneal epithelium. The finding of corneal epithelium cells in our dataset comes as a surprise, considering that corneal epithelial tissues are derived from ectoderm and hence, epidermal tissues. Since our data is based on transcriptomes from GFP+ cells isolated from eyes of Tg[foxc1b:GFP] embryos, and Foxc1b is a well-established marker of the periocular mesenchyme and ASM, we did not expect to detect corneal epithelium cells (24, 39, 64). POM and ASM cells are thought to be involved in forming anterior tissues such as the corneal endothelium, corneal stroma, the trabecular meshwork, ciliary body and iris stroma, but not the corneal epithelium. There are two possible explanations for this contradictory finding. Either, a subpopulation of corneal epithelial cells could indeed originate from neural crest cells, or this is an artefact of our transgenic line, which marks at least some corneal epithelial cells. Since ours is the first comprehensive molecular examination of POM/ASM lineage tracing, perhaps a previously unidentified population of POM does differentiate and incorporate into corneal epithelium. Support for this hypothesis stems from the fact that even at 48hpf, we detect a significant number of epithelial like cells. However, we consider the latter possibility of CEp cells originating as an artifact of the transgenic line more likely. Perhaps there is leaky expression from the transgene in corneal epithelium, as we do observe GFP+ cells, particularly at older stages, anterior of the lens (Fig 1A). Yet, at this point we cannot confidently rule out either possibility. Of the cells we identify as corneal epithelium they are represented by two clusters CEp1 and CEp2. Furthermore, determining if these clusters in fact represent two cell types is difficult. On the one hand, several marker genes such as *krt5* can be found in both clusters at the early time points but on the other, there appear to be functional differences between CEp1 and CEp2. While CEp1 mostly expresses keratins and collagens, as would be expected for this tissue, we found several markers of the immune system and stress response in CEp2, such as *anxa1a, b, c* and *serp1* (29, 65). Accordingly, CEp2 might represent a corneal immune cell type, as has been described in other organisms (66).

Corneal endothelium cells were observed to associate with the corneal stroma since both cell types closely intertwined and expressing very similar genes during early stages. In fact, we were unable to identify an independent cluster of corneal stroma cells in any of the analyzed time points. This comes as no surprise, since the stroma is known to be populated with a second wave of POM cells, after an initial wave forms the foundation of the endothelium (10). We found one cluster representing corneal endothelium and stroma, though it diverged into two clusters at 144hpf. Additionally, we found another cluster of the corneal endothelium in combination with the annular ligament cluster AL2. The major difference between the two is that the former strongly expresses markers such as *pmp22a* and *dcn*, the latter was found to also upregulate the expression of annular ligament markers such as *scinla* and *hgd*. In fact, a clear separation of annular ligament and endothelial markers at these early stages is challenging. Particularly, the annular ligament makers are broadly expressed and due to the proximity of these tissues, a clear-cut separation may not be possible at the early developmental stages. In addition to AL2 we also found another group of annular ligament cells named AL1, that separates into multiple clusters at later time points. Stratification of AL1 may represent differentiation of drainage structures analogous to the Schlemm’s canal. While the markers for both, AL1 and AL2 often overlap in expression, there are key differences in gene expression between the two cell types. For instance, the known TM and glaucoma marker *moyc* can only be found in AL2 (28, 67), just like the marker gene *lxn*, a homolog of *rarres1*, which is associated with the regulation of the fatty acid metabolism (18, 68). AL1 on the other hand strongly expresses genes involved in general development like *mcm7* and *pcna* (69–71). However, during this early time of development, analysis of functional differences remains highly speculative since the cells are not yet fully differentiated. Nonetheless, finding potentially different cell types within the annular at the early stages is exciting, since this provides a first insight into the mechanisms governing specification and assembly of this tissue.

### Trajectory of corneal and annular ligament cells throughout development

By collecting our data at multiple time points throughout early development, we were able to track cellular trajectories over time. To do so, we first manually compared gene expression of all clusters to determine if a potential relation between individual clusters within or among different time points exists. Subsequently, we performed pseudotime analysis of all combined time points, and analyzed variation in gene expression throughout the established trajectories. Manual comparison revealed that both clusters of the corneal epithelium not only share a common origin, but also are distinct from all remaining clusters, including the corneal endothelium and stroma. We found a close relation between the annular ligament and the corneal endothelium and corneal stroma clusters, indicating a common developmental origin of these tissues. When we compare our results to what is known about the physiology of AS development, they can be explained as follows. The corneal epithelium originates from epidermal cells, whereas corneal endothelium, stroma and annular ligament originate from POM cells, or more precisely ASM cells (39, 72, 73). The corneal epithelium begins development as epidermal cells overlaying the lens, starting at approximately 16hpf and is involved in initiating lens development. As mentioned previously, it remains peculiar that our data set identified corneal epithelium signatures since it is solely ASM derived. POM cells are well known for their migratory behavior and populate the anterior segment as ASM in different waves in chick, mouse and humans (37, 74, 75). The first wave of POM cells arriving at the anterior segment, migrates into the space between lens and corneal epithelium to form the endothelium. After this initial formation of the endothelium, a second wave arrives to migrate into the stroma. It is not clear at this point if the multiwave migration of POM and ASM is recapitulated in zebrafish. Furthermore, based on our data it may be that zebrafish derive all of the aforementioned tissues from a common progenitor pool that arrives early in development. Interestingly, recent work in our lab has observed the earliest ASM cells arriving as early as 16hpf, while the bulk of ASM are observed to arrive approximately 26-28hpf (39, 76). The results of our current study suggest that annular ligament, as well as corneal endothelium and the cells in the corneal stroma all share a common origin, a subgroup of ASM that are at part of the foxc1b derived lineage. We have previously shown that ASM cells are a heterogenous population during AS colonization, so the multipotent aspect of *foxc1b* derived ASM may be lineage specific rather than universal amongst all ASM. As such, in zebrafish we can conclude that corneal stroma, endothelium and annular ligament all develop from a common progenitor. Physiologically this also makes sense as stroma, endothelium and the AL share close physical association in context of AS organization. Our pseudotime analysis corroborates our manual cluster trajectories as it shows three major trajectories within our dataset (Fig 6A). The first trajectory entails corneal epithelial cells and further indicates the common origin of CEp1 and CEp2 (Fig 6B). The second trajectory contains mainly AL1 cells, whereas the third includes both AL1 and AL2, as well as CEn and CSt, further validating a relation between these tissues. Markers for AL2 such as *scinla, lxn* or *pdgfrl* are mostly found in the third pseudotime trajectory, but not the second (Fig 6B, Fig S8). Since we can find AL1 markers such as *mcm7* and *hmgb2b* in both trajectories (Fig 6B, S8) it seems possible that there is also a relation between these groups. Migrating POM cells receive developmental signals early (74, 77) and it appears possible that if even younger stages would have been included in this study, the separate trajectories could have joined into one. Accordingly, these results provide support for the hypothesis, that there is a common origin of the corneal endothelium/ stroma and at least parts of the annular ligament.

### Conservation of genetic identity between the annular ligament and trabecular meshwork

As stated above, prior to this study, our knowledge about the gene expression profile of the zebrafish anterior segment and particularly the annular ligament was very limited. As such, our study provides the first and detailed transcriptomic atlas of the developing zebrafish AS. One of the important aspects of such data is its use for comparative studies. Zebrafish, particularly their embryos, offer a highly versatile model system for modeling human disease and high throughput for functional assays related to developing treatments or interventions. While already accepted as a suitable model for ASD, zebrafish had been seldom used to assess TM biology and therefore predisposition to glaucoma. Having generated said atlas, we were able to not only uncover genes likely involved in AL development, but also compare their expression to mammalian and importantly human TM tissues. In doing so we confirmed over 31 genes to be expressed in both the TM and AL. This is not an exhaustive list, in fact many more genes should be examined for spatiotemporal expression in zebrafish and expression in TM tissues. However, clearly there is a high degree of molecular similarity between the AL and TM therefore supporting the idea of modeling TM related ASD and glaucoma pathologies. In addition to genes previously shown to associate with TM, we also identified several novel genes that are either AL specific or are not expressed in fully developed TM. Here we want to point out five of those genes previously not associated with the AS, that stood out due to their strong gene expression in our *in situ* hybridization experiments. *Hgd* and *phgdh* both are part of the pathway of tyrosine and phenylanaline degradation and we found them both to be strongly expressed in the AL and CEn tissues (see Fig 9). Mutations of *hgd* have already been proven to be the cause of a disease named Alkaptonuria, which can also affect the eye (78, 79). However, their function in the anterior segment is, to our knowledge, not known. *Stmn1a* is involved in microtubule organization and it’s homologs have previously been associated with macular degeneration of the retina (80). However, we found it to be strongly expressed in AL and CEn tissues (Fig 9). Also, when projected onto publicly available single cell transcriptomes of human tissues of the anterior segment, we found it to be widely expressed, pointing to a potentially broader function also in these tissues. *Cndp1*, a metalloproteinase family member, that has been associated with diabetic nephropathy shows a very specific staining in our study in the annular ligament (Fig 9) (81). Organization and likely remodeling of the extracellular matrix are important for TM function, hence, it may be worthwhile to determine the role that cndp1 plays in these processes. Finally, *si:211-251b21.1* is a predicted kainate receptor. Kainate receptors are known to play important roles in different tissues, including the retina and the cornea (82, 83). Due to its highly enriched expression in the AL (see Fig 9A), we assume that this particular gene might play an important function for the AL. Future functional studies will need to be carried out to assess the functional contribution of these genes to AL development as well as association with ASD or TM deficiencies.

### Zebrafish as a bona fide model for anterior segment disease

The use of zebrafish as a model for different developmental processes is well established. However, some previous studies have postulated that zebrafish should only be used with care, when applying it as a model for ocular diseases, affecting the TM or cornea due to differences in structure and molecular makeup (10, 26, 84). On the contrary, other studies suggested that zebrafish could indeed be used as a great model for certain ocular diseases and in fact started using it in this respect (8, 20–25). With this study we aim to provide a base for a better understanding of the anterior segment with focus on the annular ligament, and highlight the molecular connection it shares with the TM at early developmental time points. There are undoubtedly differences in morphology and molecular makeup of the zebrafish tissue, compared to the human TM (10, 23, 26, 84). However, having found that many TM markers described in human tissue are expressed in the AL, we strongly believe that both tissues share developmental and molecular pathways. This hopefully encourages the use of zebrafish as a functional model for disease related to TM function, such as glaucoma (4, 7).

## Acknowledgements

We thank Dr. Jeramiah Smith for invaluable assistance with Monocle3 and cell ranger data analysis. This study was funded by NIH-NEI R01EY027805-01A1 and funding from the Knights Templar Eye Foundation for OV.

## SUPPLEMENTAL FIGURES

**Figure S1**: <u>Marker gene expression distribution in AS associated clusters at 48hpf</u>. Heatmap distribution of canonical AL1 (hmgn2, hmgb2b, mcm7, pcna), AL2 (scinla, lxn), CEp1 (krt5, krt91, epcam), CEp2 (krt4, icn2, cyt1), CEn (pmp22a, ctnnb2) or CSt (dcn, lum) markers for AS associated clusters at 48hpf.

**Figure S2**: <u>Marker gene expression distribution in AS associated clusters at 72hpf</u>. Heatmap distribution of canonical AL1 (hmgn2, hmgb2b, mcm7, pcn1), AL2 (scinla, lxn), CEp1 (krt5, krt91, epcam), CEp2 (krt4, icn2, cyt1), CEn (pmp22a, ctnnb2) or CSt (dcn, lum) markers for AS associated clusters at 72hpf.

**Figure S3**: <u>Marker gene expression distribution in AS associated clusters at 96hpf</u>. Heatmap distribution of canonical AL1 (hmgn2, hmgb2b, mcm7, pcna), AL2 (scinla, lxn), CEp1 (krt5, krt91, epcam), CEp2 (krt4, icn2, cyt1), CEn (pmp22a, ctnnb2) or CSt (dcn, lum) markers for AS associated clusters at 96hpf.

**Figure S4**: <u>Marker gene expression distribution in AS associated clusters at 120hpf</u>. Heatmap distribution of canonical AL1 (hmgn2, hmgb2b, mcm7, pcn1), AL2 (scinla, lxn), CEp1 (krt5, krt91, epcam), CEp2 (krt4, icn2, cyt1), CEn (pmp22a, ctnnb2) or CSt (dcn, lum) markers for AS associated clusters at 120hpf.

**Figure S5**: <u>Marker gene expression distribution in AS associated clusters at 144hpf</u>. Heatmap distribution of canonical AL1 (hmgn2, hmgb2b, mcm7, pcna), AL2 (scinla, lxn), CEp1 (krt5, krt91, epcam), CEp2 (krt4, icn2, cyt1), CEn (pmp22a, ctnnb2) or CSt (dcn, lum) markers for AS associated clusters at 144hpf.

**Figure S6:** Corneal and annular ligament marker gene expression over time. **A)** Average mean expression measurements from CEp1 clusters for genes *krt5 and krt91* over developmental time. **B)** Average mean expression measurements from CEp2 clusters for *krt4* and *icn2* over developmental time. **C)** Average mean expression measurements from AL1 clusters for genes *myoc, hmgn2* and *hmgb2b* over developmental time. **D)** Average mean expression measurements from AL2+CEn clusters for genes *scinla, ctnnb2, pmp22a* and *lxn* over developmental time. **E)** Average mean expression measurements from CEn clusters for genes *ctnnb2* and *pmp22a* over developmental time. **F)** Average mean expression measurements from CEn+CSt clusters for genes *dcn, lum, ctnnb2* and *pmp22a* over developmental time.

**Figure S7:** Verification of Pigment cell expression patterns. WISH expression analysis of select pigment cell associated genes at 24, 72 and 120hpf displayed in dorsal and lateral views.

**Figure S8:** <u>Gene expression distribution in the pseudotime analysis of the combined datasets</u>. Heatmap distribution of (A) sparc, lgals1l1, (B) col1a2, col4a5, col1a1b, (C) anxa1c, pnp5a, (D) fabp11a, (E) hgd, si:ch211-251b21.1 and (F) fmoda projected onto the dataset, including the aggregated data of 48hpf, 72hpf, 96hpf, 120hpf and 144hpf.

**Figure S9:** <u>Conserved and novel CEp associated gene expression patterns</u>. **A)** WISH expression analysis of mammalian corneal epithelium conserved CEp associated genes at 24, 72 and 120hpf displayed in dorsal and lateral views. **B)** WISH expression analysis of potentially novel CEp associated genes at 24, 72 and 120hpf displayed in dorsal and lateral views.

**Figure S10:** <u>Conserved and novel AL associated gene expression patterns</u>. **A)** WISH expression analysis of mammalian TM conserved AL associated genes at 24, 72 and 120hpf displayed in dorsal and lateral views. **B)** WISH expression analysis of potentially novel AL associated genes at 24, 72 and 120hpf displayed in dorsal and lateral views.

