## Supplementary Figures 1-10 for "Temporal single cell transcriptome atlas of zebrafish anterior segment development reveals high degree of conservation between the trabecular meshwork and the annular ligament": Manuscript SCT Supplementry Figures.pdf

### Supplemental Figure 1

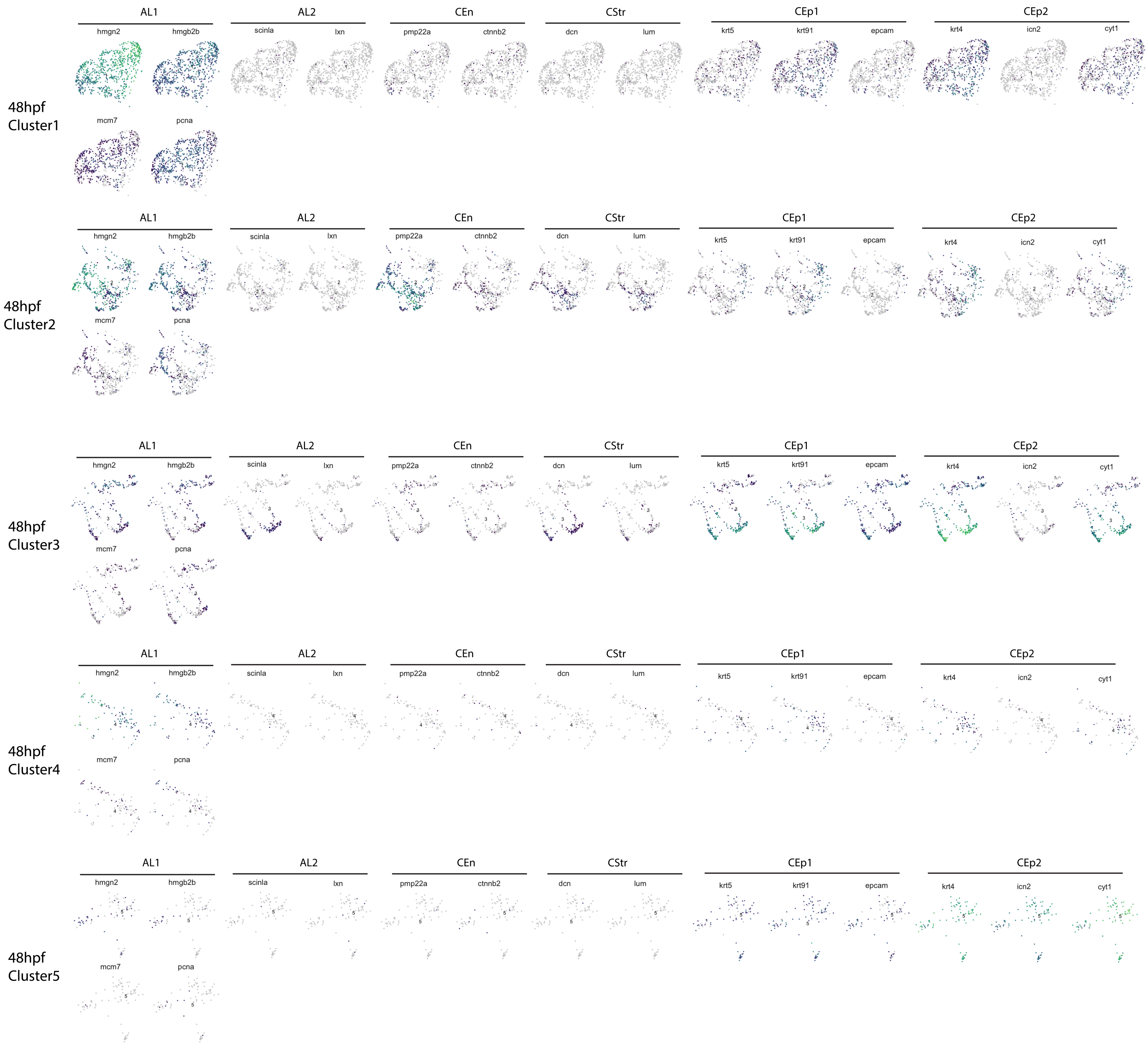

### Supplemental Figure 2

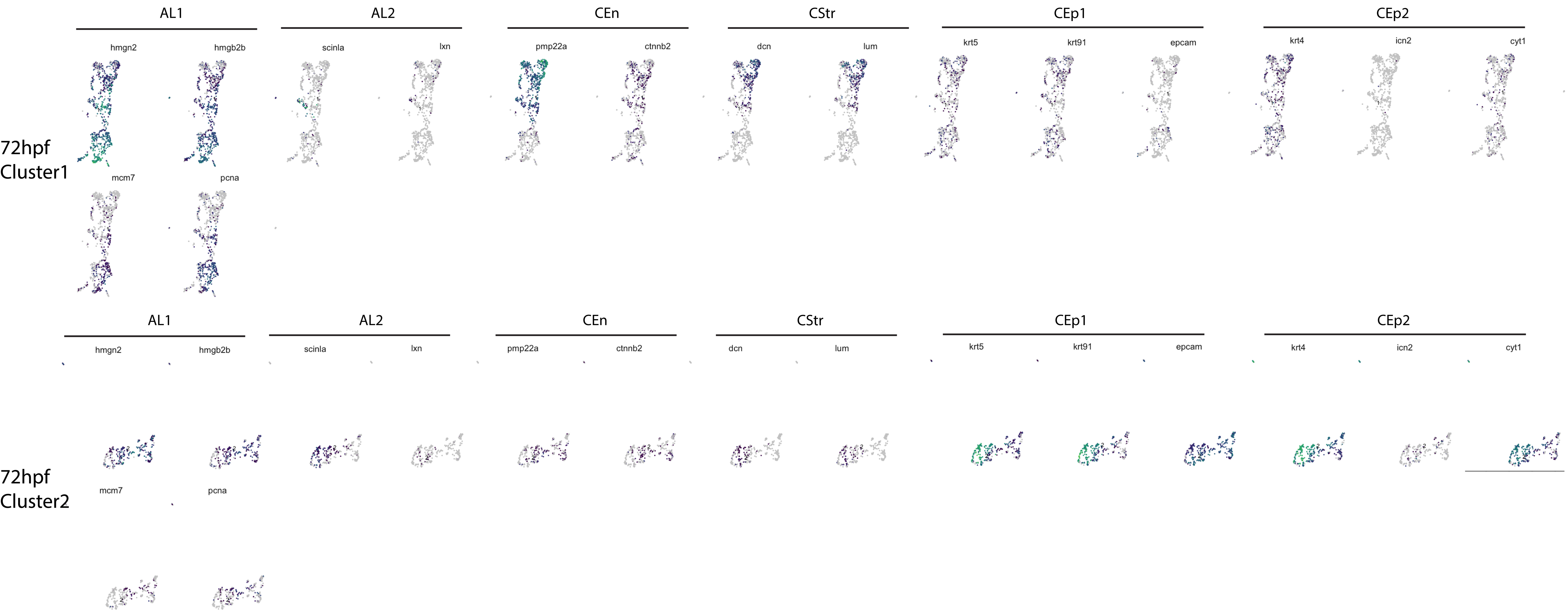

### Supplemental Figure 3

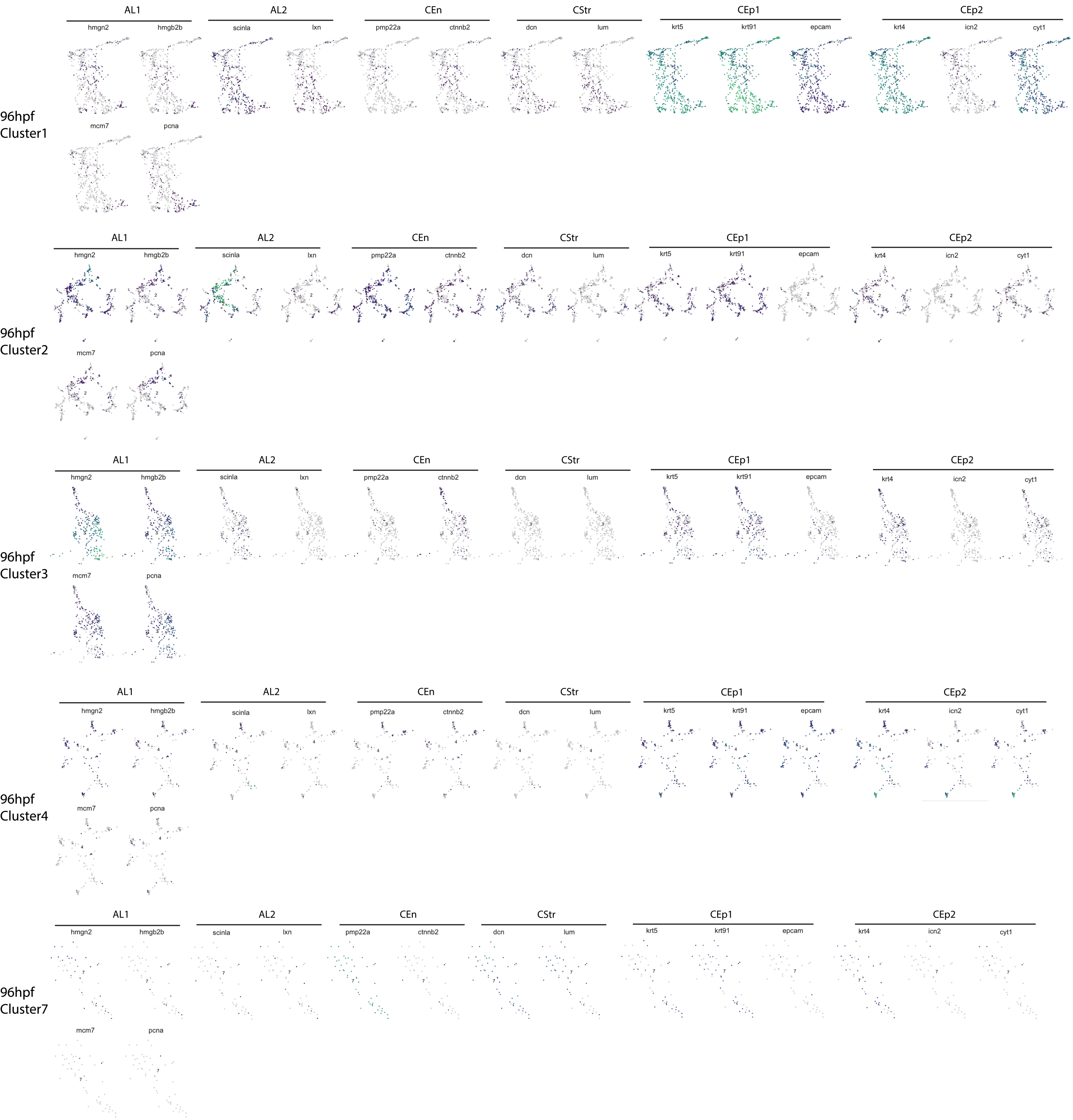

### Supplemental Figure 4

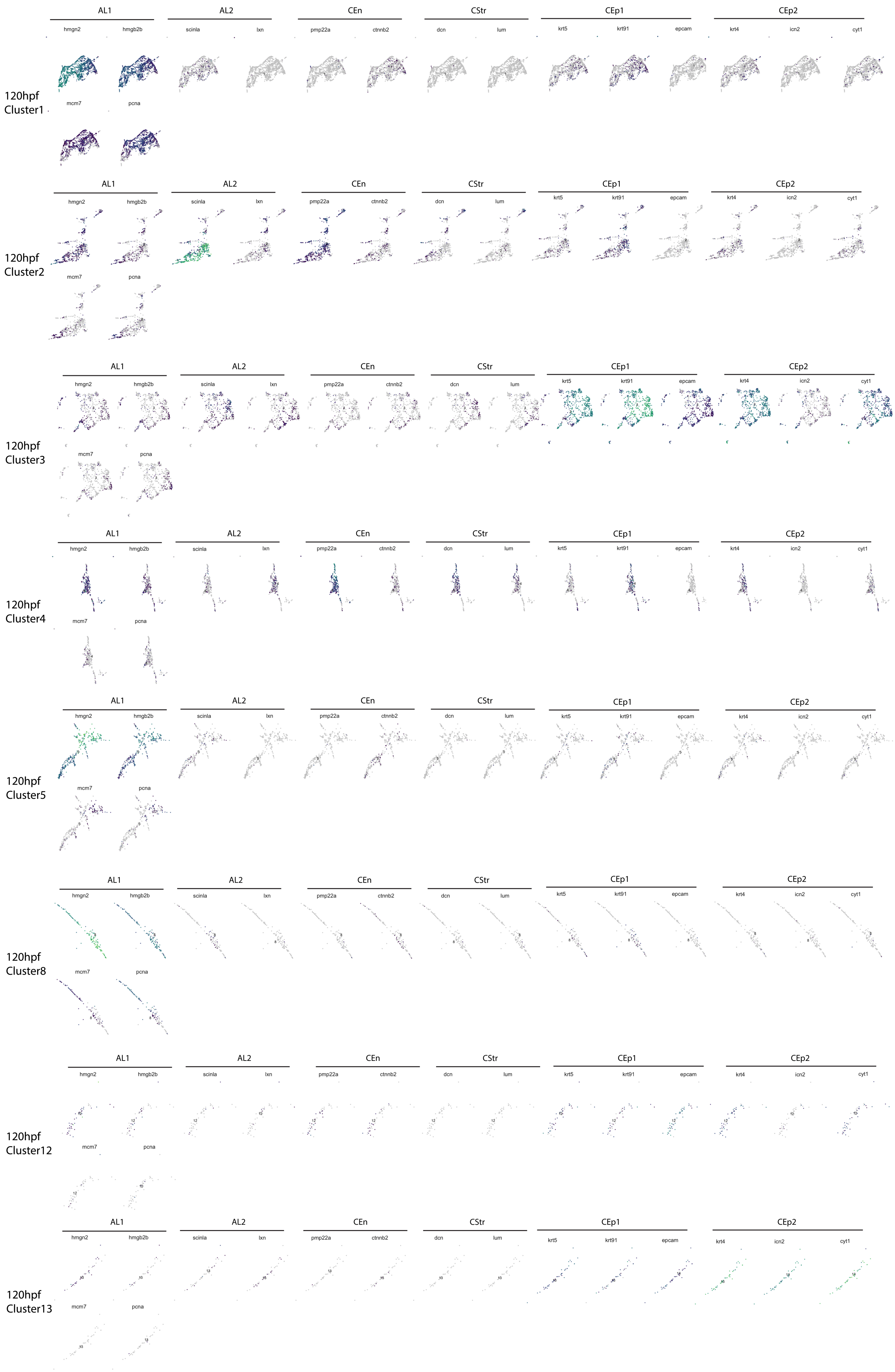

### Supplemental Figure 5

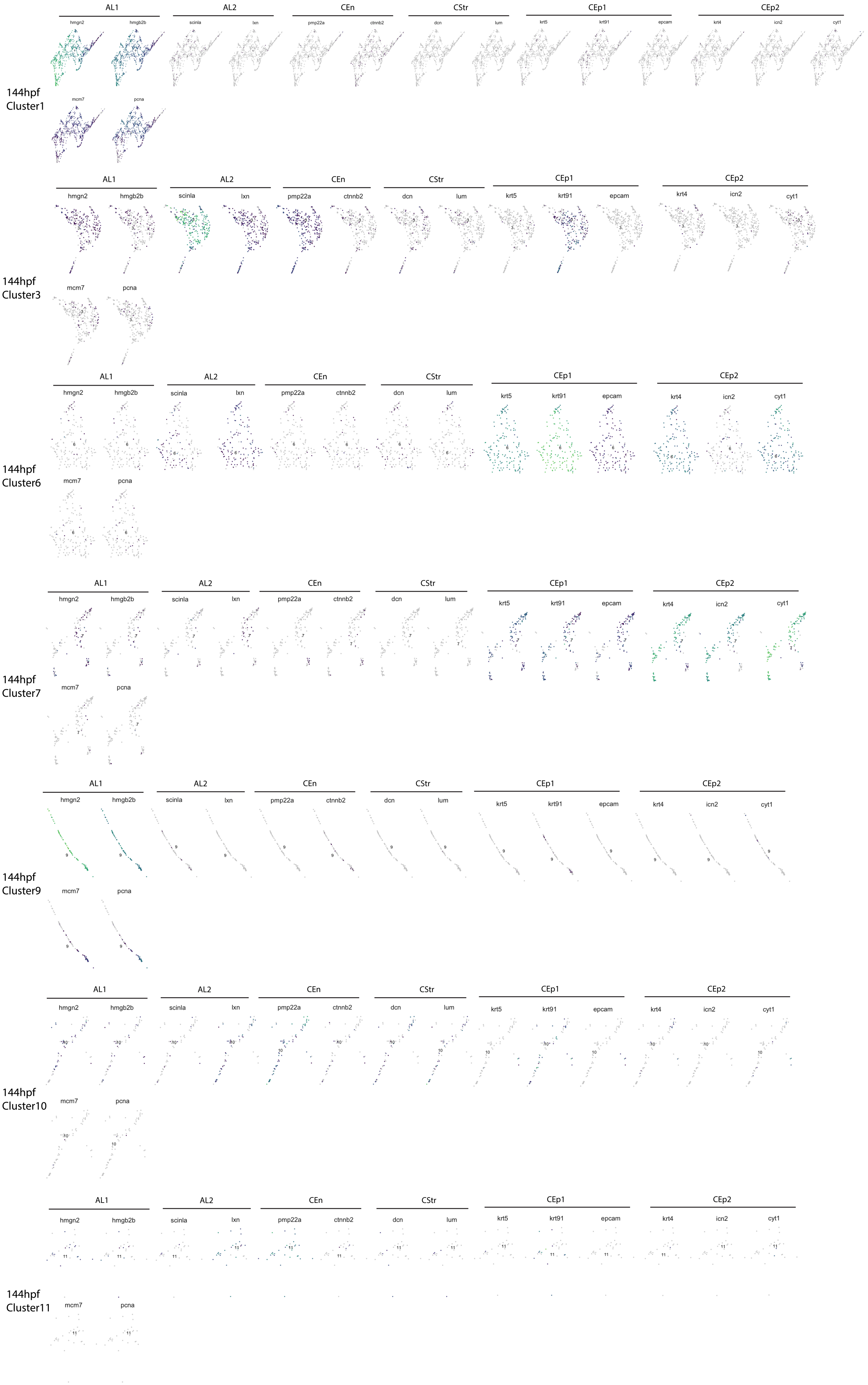

### Supplemental Figure 5

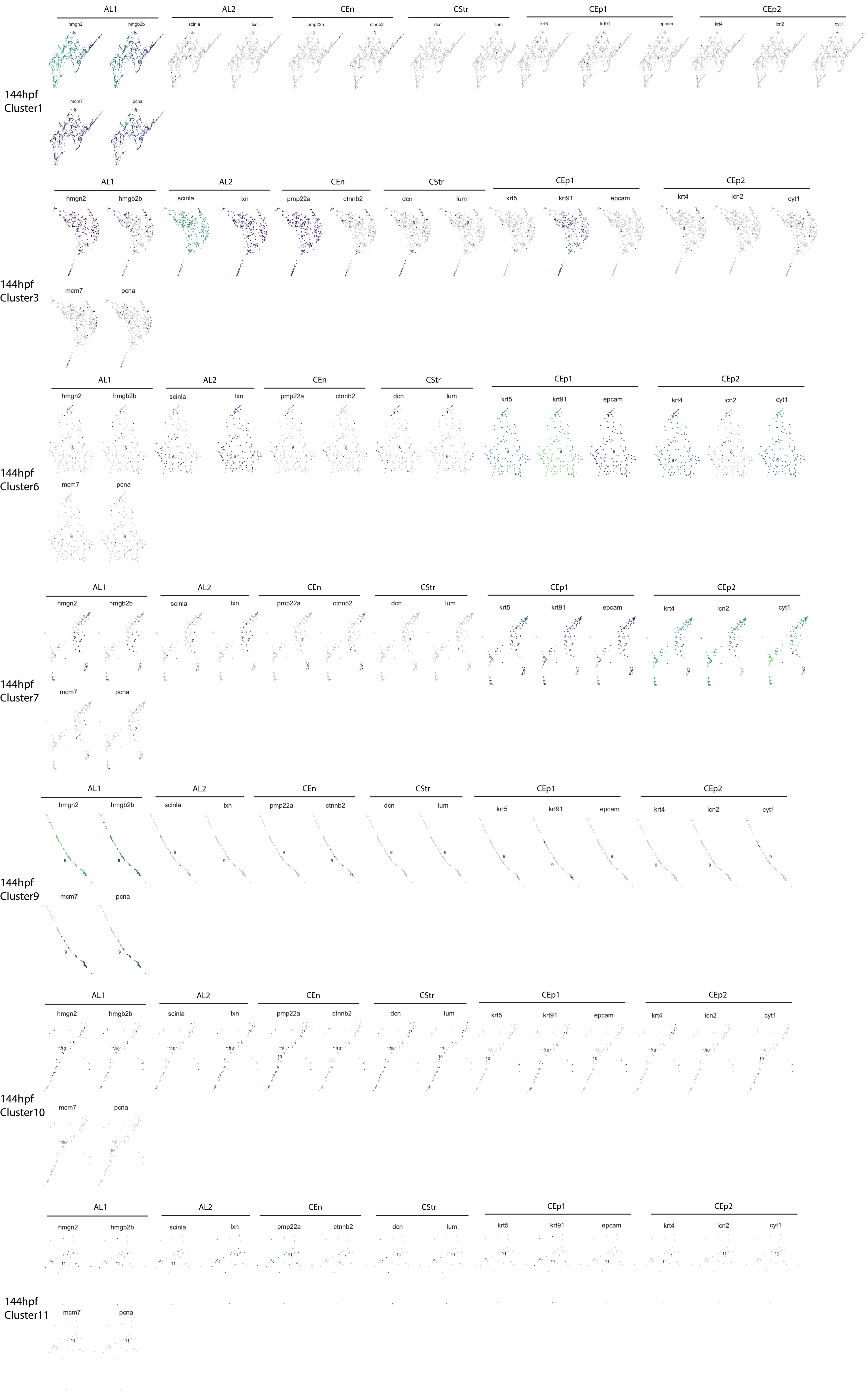

### Supplemental Figure 6

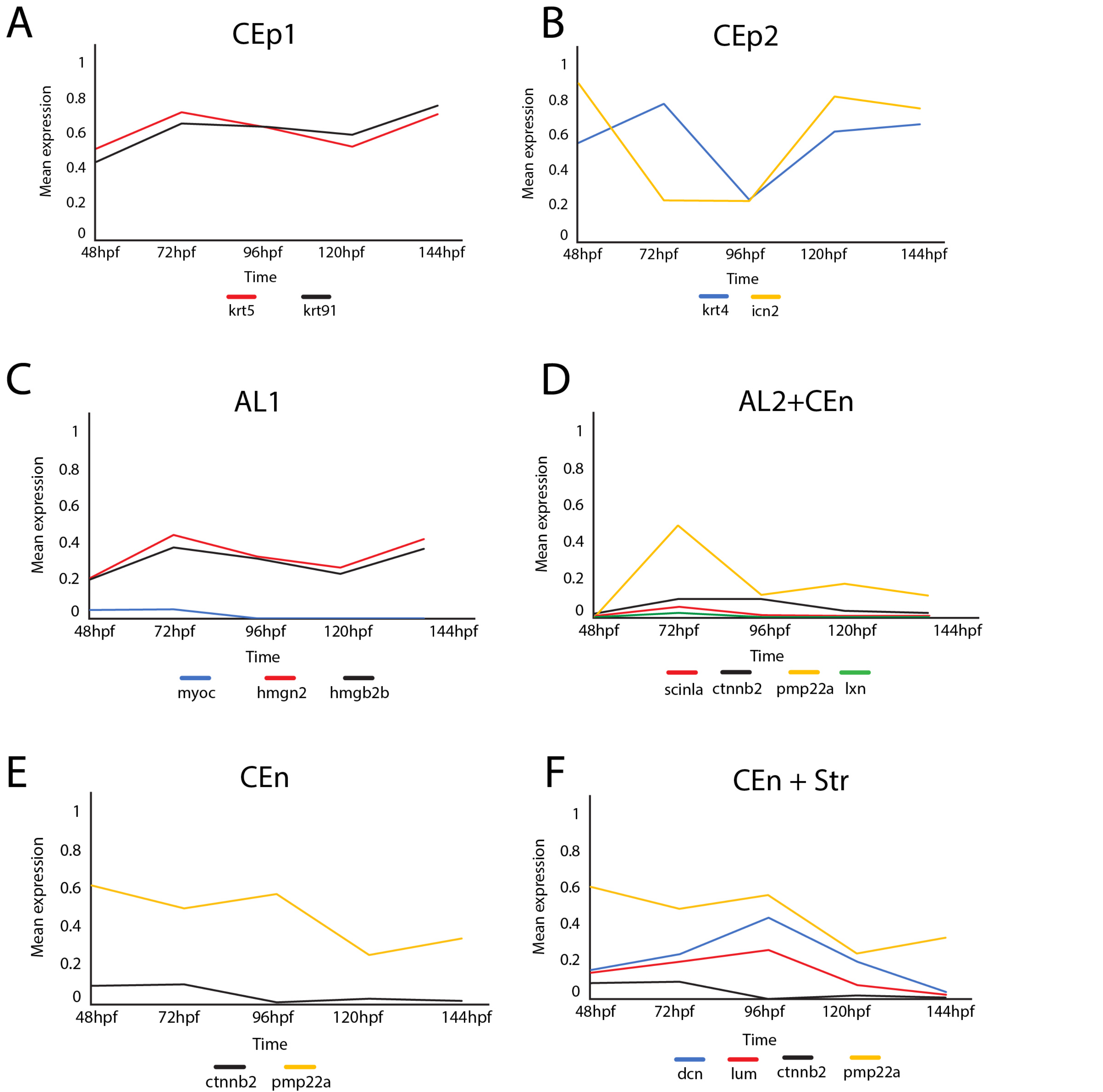

### Supplementary Figure 7

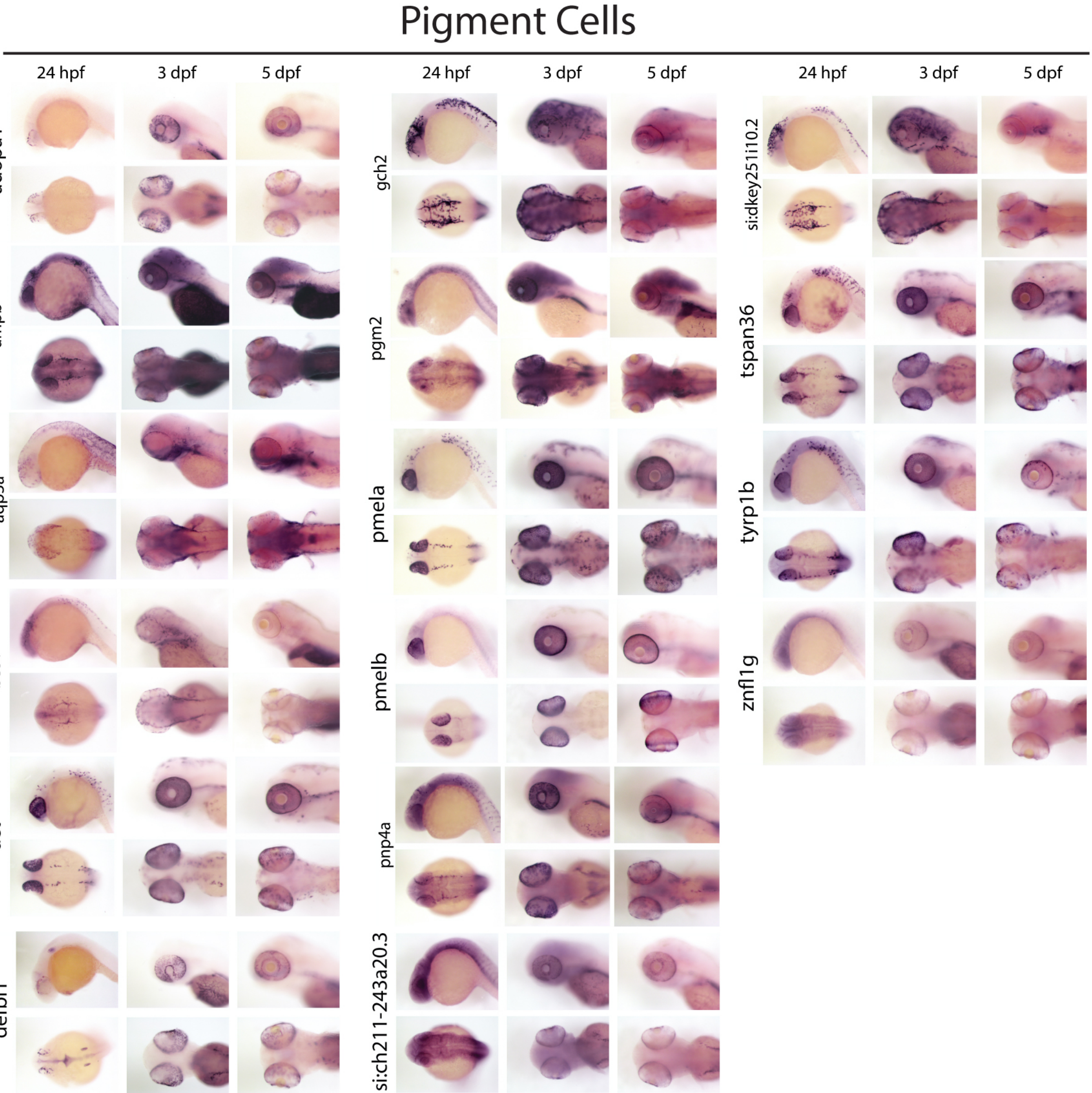

### Supplemental Figure 8

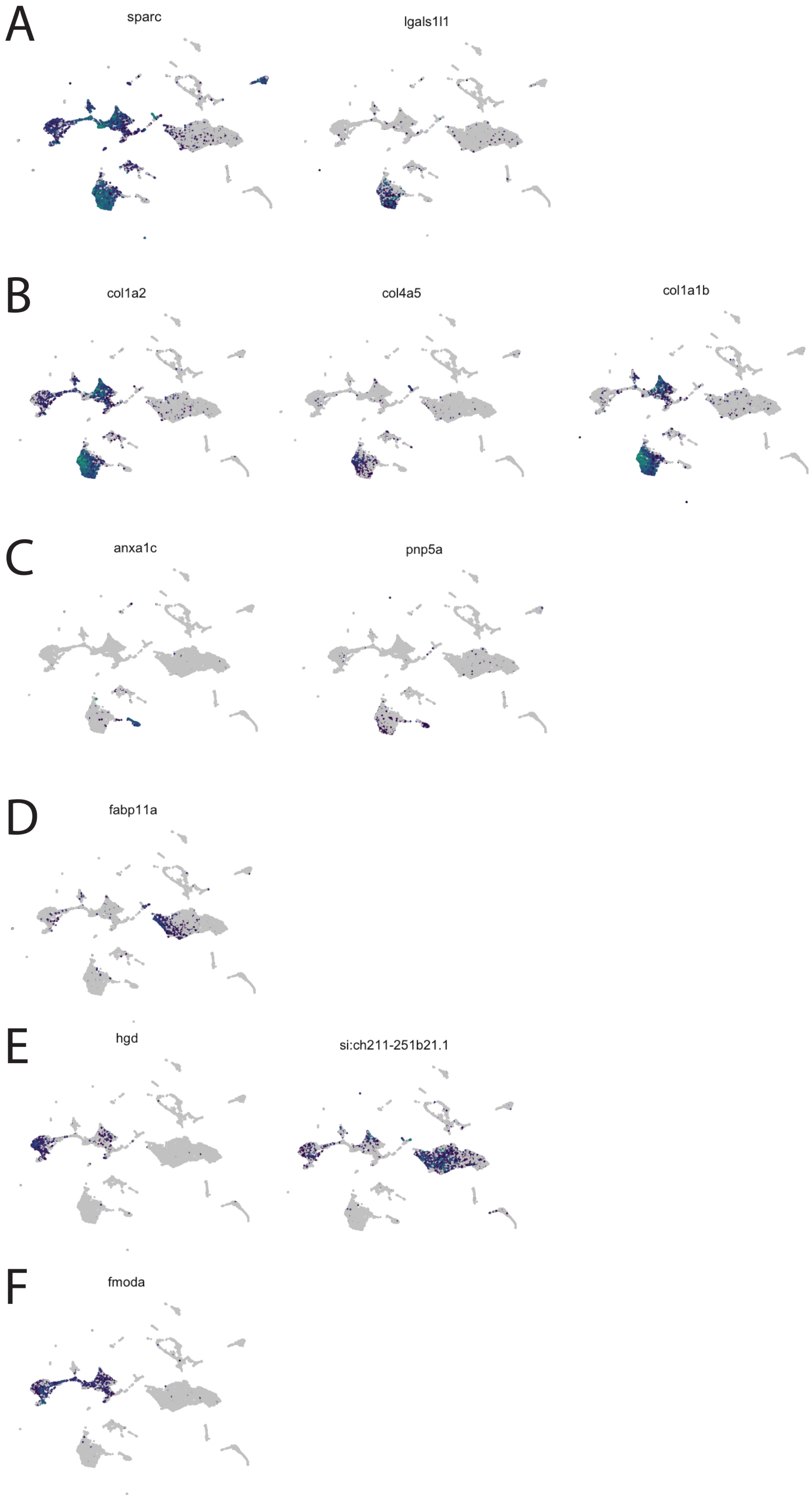

### Supplemental Figure 9

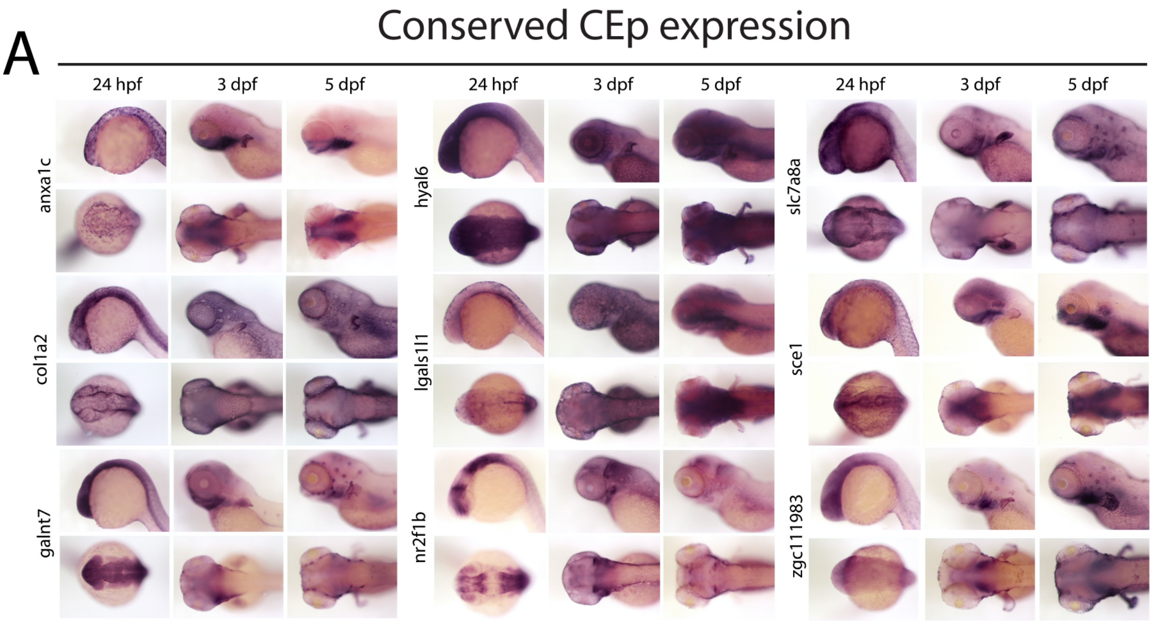


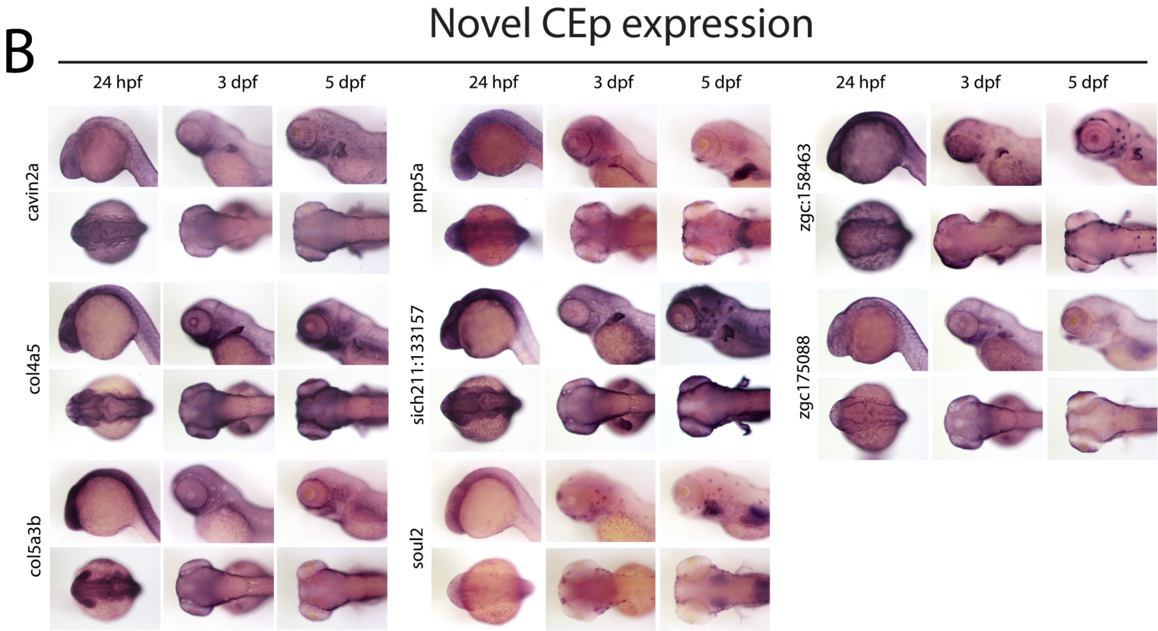

### Supplemental Figure 10

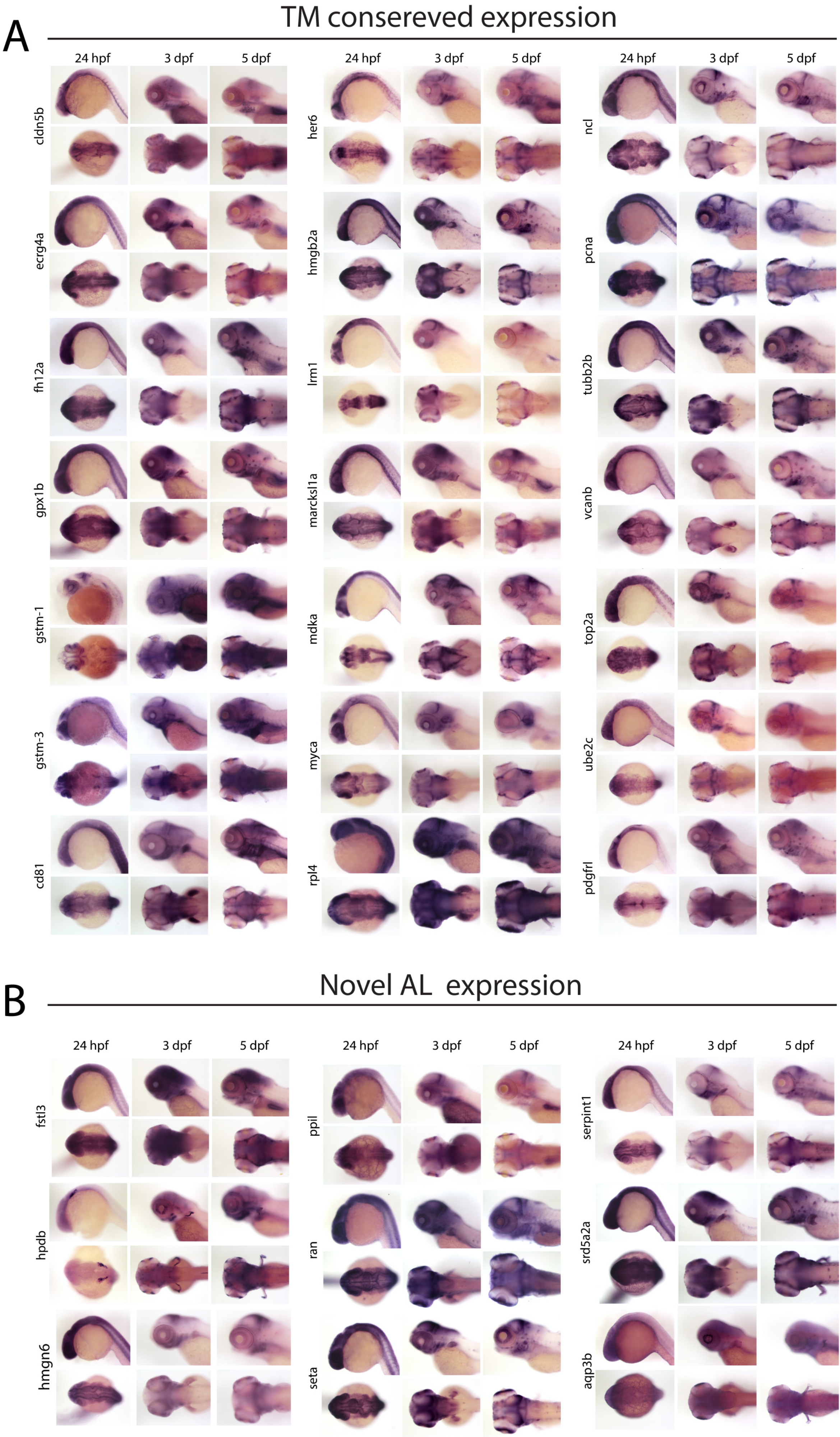
